# Cryo-EM reveals disrupted human p97 allosteric activation by disease mutations and inhibitor binding

**DOI:** 10.1101/2021.08.10.455711

**Authors:** Brian Caffrey, Xing Zhu, Alison Berezuk, Katharine Tuttle, Sagar Chittori, Sriram Subramaniam

## Abstract

The human AAA ATPase p97, a potential cancer target, plays a vital role in clearing misfolded proteins. p97 dysfunction is also known to play a crucial role in several neurodegenerative disorders. Here, we present cryo-EM structural analyses of four disease mutants p97^R155H^, p97^R191Q^, p97^A232E^, p97^D592N^, as well as p97^E470D^, implicated in resistance to the drug CB-5083. These structures demonstrate that the mutations affect nucleotide-driven p97 allosteric activation by predominantly interfering with either the coupling between the D1 and N-terminal domains (p97^R155H^ and p97^R191Q^), the inter-protomer interactions (p97^A232E^), or the coupling between D1 and D2 nucleotide domains (p97^D592N^, p97^E470D^). We also show that binding of the competitive inhibitor CB-5083 to the D2 domain prevents conformational changes similar to that seen for mutations that affect coupling between D1 and D2 domains. Our studies enable tracing of the path of allosteric activation across p97 and establish a common mechanistic link between active site inhibition and defects in allosteric activation by disease-causing mutations.

## INTRODUCTION

p97 is a member of the classic clade of AAA+ ATPases and is an essential cellular enzyme. p97 is composed of six protomers forming a homo-hexamer with an N-terminal domain (NTD) and two tandem ATPase domains D1 and D2 (1). p97 is an essential protein in regulating cellular homeostasis from membrane fusion, Endoplasmic Reticulum-Associated Degradation (ERAD), Mitochondrial-Associated Degradation (MAD), Chromatin-Associated Degradation to NF-κB activation (2). A common thread among these applications is the recruitment of numerous cofactors to process ubiquitinated substrates, through the generation of mechanical energy from ATP hydrolysis. Critical functions for p97 include the translocation and restructuring of proteins from large cellular structures such as organelle membranes and extraction of ubiquitinated client proteins to facilitate their degradation through the ubiquitin-proteasome system. In this role, p97 recruits ubiquitin-binding cofactors to denature ubiquitinated substrates by pulling the polypeptide through the central pore (Cooney et al., 2019; Twomey et al., 2019).

Cryo-EM analyses of full-length p97 in the substrate-free form have established the overall organization of N, D1 and D2 domains (Fig. 1A) in three distinct quaternary conformations that correspond to distinct nucleotide occupancies of the D1 and D2 nucleotide-binding sites (Fig. 1B)(5). Conformation I is observed when ADP is bound to both the D1 and D2 domains, while conformation II is observed when ADP in the D2 binding site is exchanged with ATPγS, which is a slowly hydrolysable ATP analog to preserve the ATP-bound structure for cryo-EM analysis. Relative to conformation I, there is a rotational twist of the D2 domain with relatively minimal changes in the D1 and N domains. Conformation III is observed when ATPγS is bound to both D1 and D2 nucleotide-binding sites and results in substantial tertiary and quaternary structural rearrangements in the D1 and N domains. The N-domain in conformation III adopts a distinct “up” position, enabling its binding to endogenous cofactors required for subsequent substrate binding and processing. The nucleotide occupancy in these three conformational states is consistent with the higher affinity of the D1 domain for both ADP and ATP ligands relative to the D2 domain, with a ∼40-fold lower K_D_ in D1 relative to D2, as measured by Surface Plasmon Resonance (SPR) experiments (6). All three conformations of substrate-free p97 display 6-fold symmetry. However, substrate-binding results in the conversion of the hexameric arrangement to a spiral arrangement of the six protomers similar to the quaternary conformation observed for most other members of the AAA+ ATPase family (Cooney et al., 2019; Twomey et al., 2019).

**Fig. 1:**
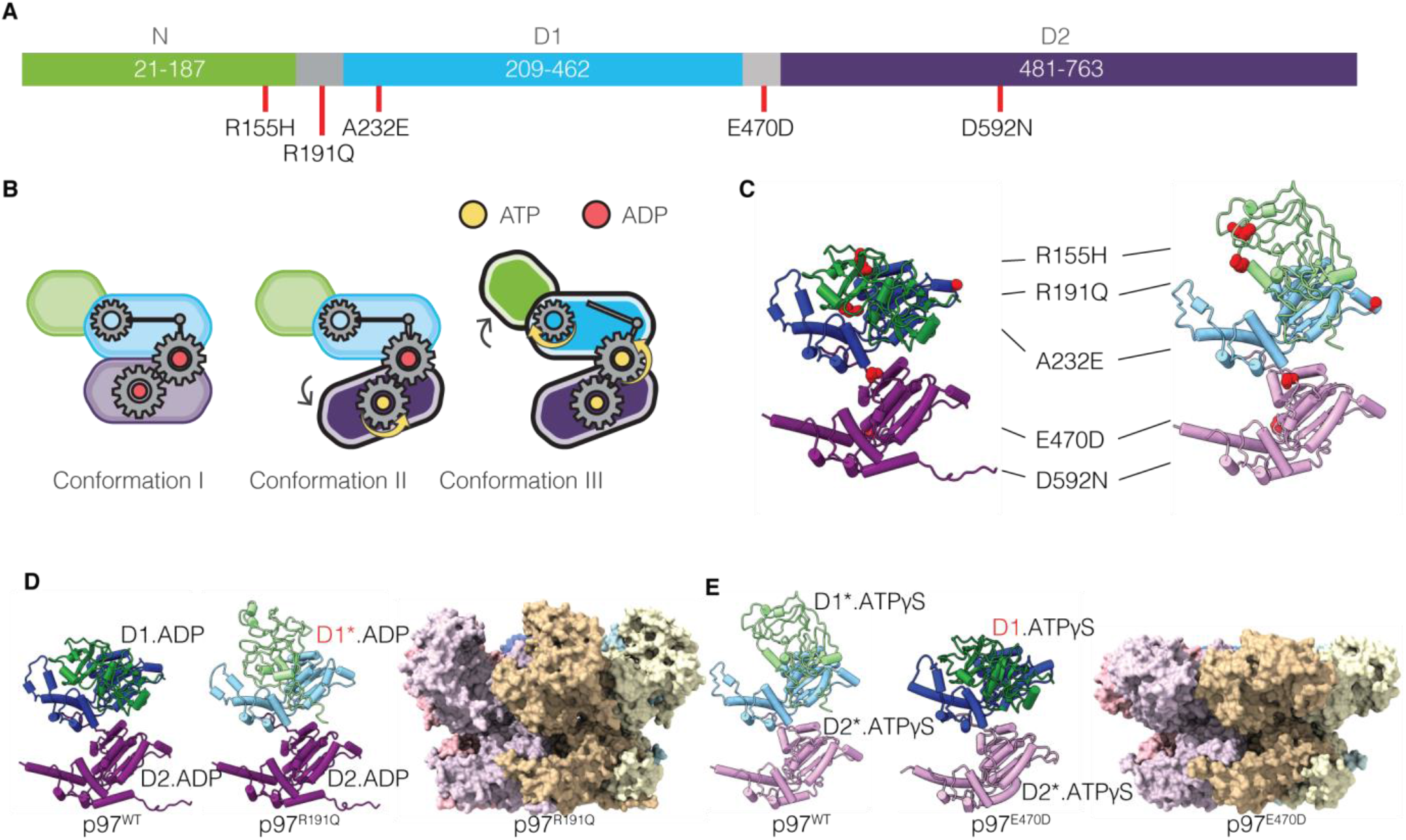
Cryo-EM Structures of p97 Disease Mutants. **(A)** Schematic of human p97 sequence identifying the location of the three p97 domains and locations of the mutations analyzed in this study. **(B)** Schematic representation of the three conformations observed for p97^WT^. **(C)** Location of mutations labeled in **(A)** in the 3D structure of a p97^WT^ protomer in ADP-bound (left panel, PDB: 5FTL) and ATP*γ*S-bound (right panel, PDB: 5FTN) states. **(D)** Atomic models for the ADP-bound p97^WT^ and p97^R191Q^ protomers (left) and space-filling model of the p97^R191Q^ hexamer in the ADP-bound state (right). **(E)** Atomic models for the ATP*γ*S-bound p97^WT^ and p97^E470D^ protomers (left) and space-filling model of the p97^E470D^ hexamer in the ATP*γ*S-bound state (right). The darker and lighter colors of the D1 and D2 domains in this and subsequent figures reflect the conformation of the domains observed in p97^WT^. For example, as shown in **(D)**, the conformation of the ADP-bound D2 domain in p97^R191Q^ the same as that observed for p97^WT^, but that of the ADP-bound D1 domain is different, and closer to that seen for the ATP*γ*S-bound D1 domain.

Due to the essential role played by p97 in cellular processes, it is no surprise that a number of multisystem diseases are associated with mutations and dysfunctions in p97. Diseases arising from protein degradation defects (7) and DNA repair (8) such as cancer, viral infections such as the poliovirus (9) and even neurodegenerative disorders all implicate p97 as crucial in disease progression. p97 is therefore an attractive therapeutic target (10). The majority of p97 mutations are associated with a rare disease called Multi System Proteinopathies-1 (MSP-1) which is characterized by a cluster of muscular, bone, and neurological illnesses known as Inclusion Body Myopathy (IBM); Paget’s Disease of the Bone (PDB); and FrontoTemporal Dementia (FTD)(11). As these mutations tend to cluster in the N- and D1-domains of the p97 protein and the majority of p97 cofactors bind the N-terminus of the protein, there is likely cofactor involvement in the mechanism of disease. These mutants are also commonly characterized by an increase in basal ATPase activity (12) and higher substrate processing relative to wild-type p97 (13), although these effects have not been shown to be directly correlated. Specific mutations in the D2 domain of p97 have also been identified in closely related neurological illnesses such as Amyotrophic Lateral Sclerosis (ALS) (14) which tend to have less pronounced muscular and bone involvement suggesting a subtle but significant difference in the mechanism of disease. Still, other mutations in the D1-D2 linker are implicated in cancer drug resistance (15), reflecting the diverse role of p97 within the cell and its importance in different tissues. A central unanswered question in understanding the origin of the disease phenotype is whether the functional consequences can be traced to mutation-induced alterations in protein structure, which can then be used to account for the broad variance and severity in the phenotype of these diseases and the development of therapeutic interventions.

Prior structural information on p97 mutants largely comes from X-ray crystallographic studies of N-D1 truncated p97 constructs (16), NMR investigations of protein dynamics of full-length p97 mutants (17), and cryo-EM analysis of full-length p97 in complex with Ufd1-Npl4 cofactors (13, 18). Here, we have examined full-length forms of five p97 mutants: three were chosen for their relatively high incidence in the p97-related neurodegenerative disorder MSP-1 (R155H, R191Q, A232E), one for its relatively unusual position in the D2 domain in p97 and its role in ALS (D592N) and one for its identification in resistance to the drug CB-5083 (E470D) (15).

Our experiments with these mutants were designed to explore the extent to which the mutations influence the conformational landscape. While p97 is amenable to structural analysis in both substrate-free forms as well as in the presence of bound protein substrates and co-factors, we chose to study substrate-free complexes since our goal is to unravel the intrinsic effects of the mutations on quaternary structure. Previous studies have shown that the primary effect of MSP-1 mutations is in the transition between conformations (I-III) rather than directly on the hydrolytic domains (6) or indeed cofactor-bound complexes (13).

We also report cryo-EM structures of full-length wild-type p97 complexed to CB-5083 in the presence of ADP and ATPγS, and describe how the binding of the inhibitor influences the overall quaternary arrangement of the N, D1, and D2 domains. Taken together, our results provide a detailed understanding of the structural mechanisms underlying the effects of the disease mutations, and an explanation of the resistance of the E470D mutation to inhibition by CB-5083.

## RESULTS

All five mutants displayed relative ATPase activities similar to previously reported biochemical studies (Fig. 1C and Supplementary Fig. 1A) (12, 15, 16). The structural effects of the mutations on p97 structure as determined by cryo-EM studies fall into two distinct categories where the differences in behavior relative to p97^WT^ are observed in the presence of bound ADP but not ATPγS (Fig. 1D) or in the presence of bound ATPγS but not ADP (Fig. 1E). These effects are described in greater detail below.

### R155H, R191Q, and A232E Mutants Primarily Affect the ADP-bound States

In the p97^R191Q^, p97^R155H^ and p97^A232E^ mutants, the primary variation in conformational outcome as compared to p97^WT^ is observed when the D1 and D2 domains are both occupied by ADP (Fig. 1D). In these mutants, the ADP-bound conformation of the D1 domain displays many of the structural signatures observed only upon ATPγS binding to p97^WT^, suggesting that these mutations shift the conformational equilibrium towards promoting ATP binding. This effect is most readily visible in p97^R191Q^, with p97^R155H^ and p97^A232E^ displaying similar effects in the D1 domain, but of a smaller magnitude (Fig. 2A-C and Supplementary Fig.1B-D and Fig. 2A).

**Fig. 2:**
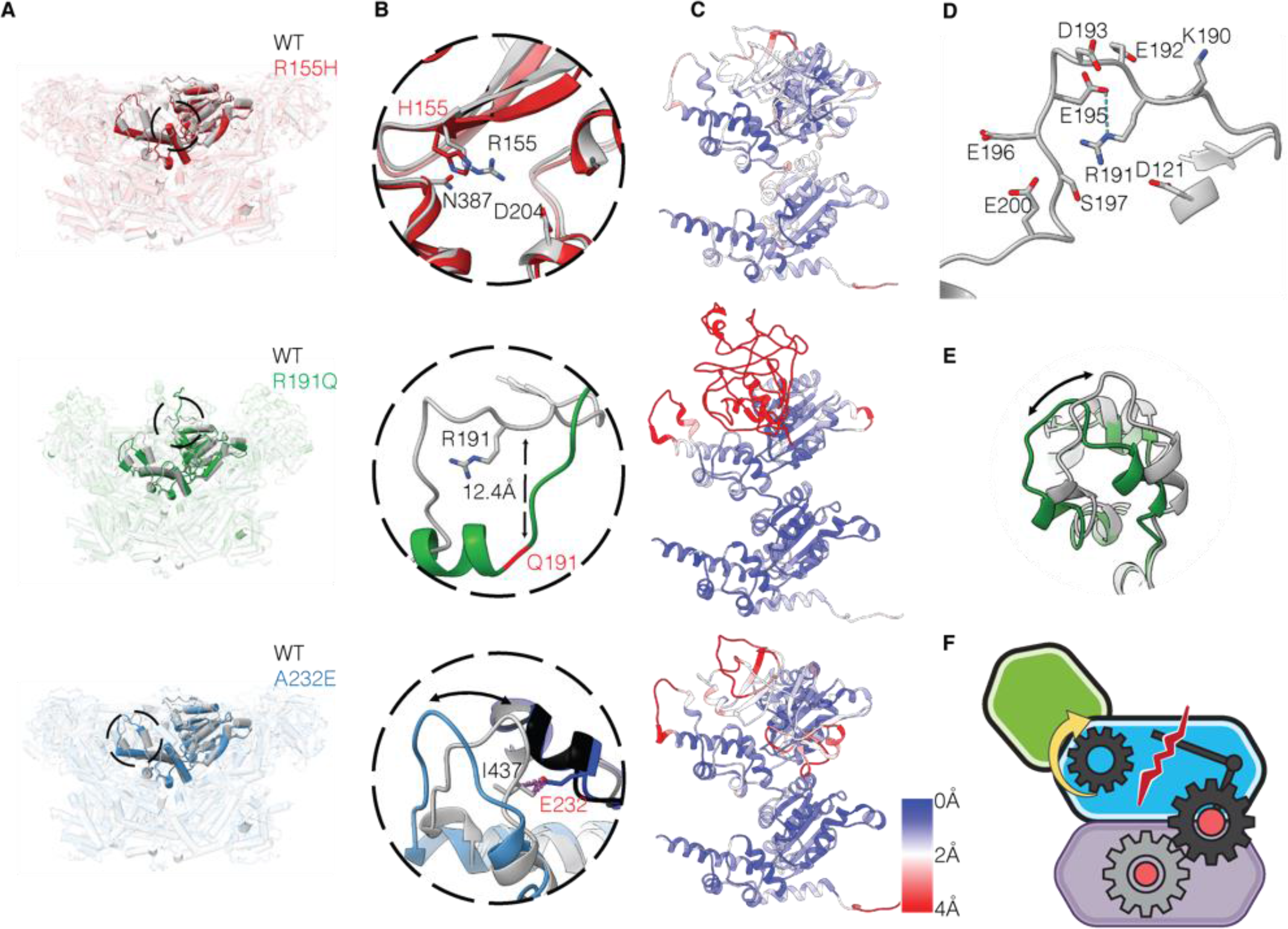
R155H, R191Q and A232E Mutations Disrupt N-D1 Domain Interactions. (**A**) **Top:** Superposition of both p97^WT^ (grey) and p97^R155H^ (red) ADP-bound hexameric structures, with the N-D1 linker and D1 domains highlighted (residues 180-460). **Middle:** Superposition of both p97^WT^ (grey) and p97^R191Q^ (green) ADP-bound hexameric structures. **Bottom:** Superposition of both p97^WT^ (grey) and p97^A232E^ (blue) ADP-bound hexameric structures. **(B) Top:** Expanded view of the site of the p97^R155H^ mutant, illustrating an increase in the interaction between residues H155 and N387 relative to WT. **Middle:** Expanded view of the site of the p97^R191Q^ mutant in the N-D1 linker region (residues 188-209), illustrating the mutant driven loop-to-helix transition. **Bottom:** Steric clash between mutant E232 and adjacent hydrophobic residue in p97^WT^ structure, modelled in chimera (van der Waals overlap of ≥0.6Å), labelled in purple. Individual Mutant D1-domains labelled in light blue/blue and WT D1-domains labelled in light grey/black. **(C) Top**: p97^R155H^ protomer is displayed showing backbone Root Mean Square Deviation (RMSD) from D1D2-ADP-bound p97^WT^ [PDB:5FTK]. **Middle:** p97^R191Q^ protomer is displayed showing backbone RMSD from D1D2-ADP-bound p97^WT^ [PDB:5FTK]. **Bottom:** p97^A232E^ protomer is displayed showing backbone RMSD from D1D2-ADP-bound p97^WT^ [PDB:5FTK]. **(D)** N-D1 linker region of ADP-bound p97^WT^ highlighting R191 and the surrounding residues, tentative hydrogen bond modelled in chimera, according to precise geometric constraints based on (20) labelled in cyan. **(E)** Expanded view of the D1 domain loop region (residues 425-445) of p97^WT^ and p97^R191Q^. **(F)** Schematic illustrating mutant protomer disruption in N-D1 interface, leading to flexibility in the N-domain conformation towards the “up” position.

R155 is located at the N-D1 interface in the ADP-bound state making contacts with polar residues from the N-D1 domain(19) (Fig.2B). Mutation to histidine may disrupt interactions resulting in greater flexibility of the N-domain, consistent with disorder in this domain, observed as a decrease in local resolution in the N-domain cryo-EM density. This finding is consistent with previously published NMR studies where the N-domain flexibility i.e., exchange between “up” and “down” conformations in the ADP-bound state, is inversely proportional to the charged/hydrogen bonding nature of the residues at position 155 (17).

R191 is located at the start of the linker connecting the N and D1 domains (Fig. 1A and Fig.2), surrounded by negatively charged residues, in particular E195 (Fig. 2B+D). R191 may stabilise the “down” position of the N-domain by maintaining an extended loop conformation in the linker. A unique aspect of the mutation-induced structural changes in the ADP-bound state of p97^R191Q^ is a dramatic ordering of the linker connecting the N and D1 domains into an *α*-helical structure between residues 190–199 (Fig. 2B and Supplementary Fig. 1B). This structural ordering is observed in p97^WT^ only when ATPγS is bound to the D1 domain, which arises from changes propagated at the other side of the linker proximal to the D1 nucleotide binding site. The dramatic shift from extended linker to an *α*-helix upon loss of charge by mutation from arginine to glutamine appear to confirm the vital role R191 plays in stabilising the “down” position in the N-domain of p97^WT^. Further inspection of the structure of p97^R191Q^ in the ADP-bound state shows a shift in the local conformation of the loop region between K425 and L445 of the D1 domain that is located at the junction between the N and D1 domains (Fig. 2E and Supplementary Fig.1B). The change in the structure of this loop is such that it is close to the conformation observed for this region in p97^WT^ upon the occupancy of the D1 nucleotide binding site with ATPγS.

Residue A232 is located at the interface between two p97 protomers, and the introduction of the glutamate at this position will result in steric repulsion (Fig. 2B) separating the putative hydrogen bonding residues T127 and T436 of the N and D1 domains of the adjacent monomer, respectively (Supplementary Fig.1C). This disruption likely accounts for the increased flexibility of the N-domain and also results in a shift similar to that of p97^R191Q^ in the loop region between K425 and L445 of the D1 domain adjacent to the E232 residue (Fig.2B and Supplementary Fig. 1D). Of note, a T127A mutation has been reported in an individual with FTD (21), further suggesting the importance of this hydrogen bonding interaction for the function of p97.

Despite varying degrees of differences in the ADP-bound state as compared to p97^WT^, all three mutants display essentially similar structures when the D1 and D2 domains are bound to ATPγS and in the state corresponding to conformation III of p97^WT^ (Supplementary Fig. 3A-D) (5). This is not surprising since R155, R191 and A232 are all exposed to solvent in conformation III, i.e. no longer interacting at the N-D1 interface, therefore the mutations at these sites are not expected to affect the quaternary conformation significantly. The effect of the ATPγS-driven conformational change is especially noteworthy in the case of the p97^R155H^ mutant because the upward movement of the N-domain results in the displacement of R155 by ∼25 Å to a solvent exposed face of the protein, away from the N-D1 interface. In the ATPγS-bound state, p97^R155H^ and other MSP-1 mutants with similar phenotypes may have an effect on cofactor or substrate binding since it is in close proximity to the site where cofactors are expected to interact with p97 (22). Indeed, the p97^R155H^ mutant has been found to have altered responses relative to p97^WT^ when bound to either p47 or p37 cofactors (23).

### E470D and D592N Mutants Primarily Affect the ATP***γ***S-bound States

In contrast to the mutants discussed above, in the p97^E470D^ and p97^D592N^ mutants, minimal structural differences are observed as compared to p97^WT^ when the D1 and D2 domains are occupied by ADP (Supplementary Fig. 4). However, when bound to ATPγS, in contrast to displaying the dramatic quaternary structural changes observed in p97^WT^, the predominant conformation displayed in these two mutants is one in which the N-terminal domain is in the “down” position in a state most closely resembling conformation II of p97^WT^ (Fig.3 and Supplementary Fig. 1E). These results imply that the mutations decouple ATPγS binding in the D1 domain from inducing the structural rearrangement of N domain from the “down” to “up” conformation.

**Fig. 3:**
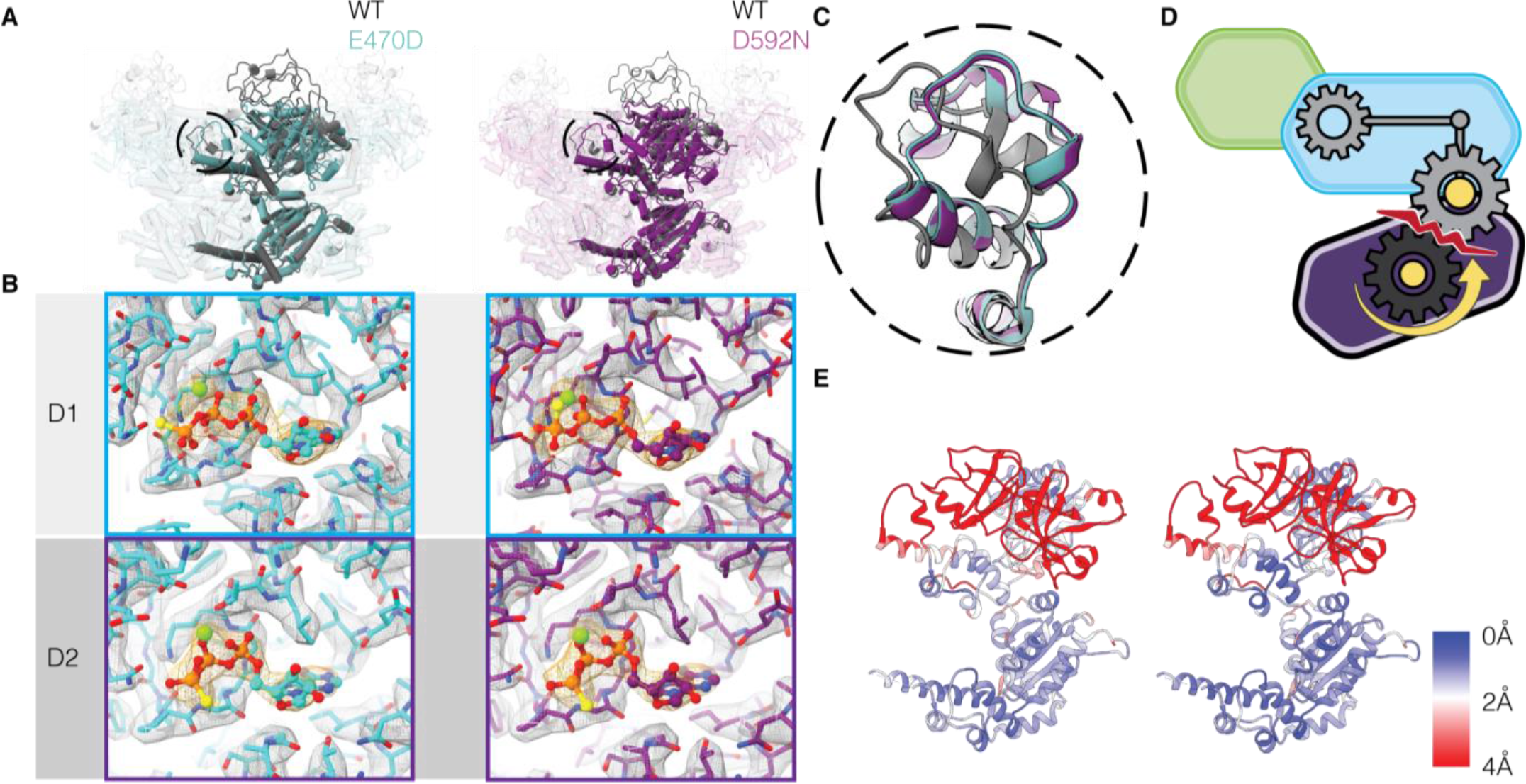
E470D and D592N Mutations Alter ATP Induced Conformational Changes. **(A) Left:** Superposition of p97^WT^ (grey) and p97^E470D^ (cyan) ATPγS-bound hexameric structures, with monomer highlighted. **Right:** Superposition of p97^WT^ (grey) and p97^D592N^ (purple) ATPγS-bound hexameric structures, with monomer highlighted. **(B)** Illustration of nucleotide occupancy in both the D1 and D2 domains of the unsharpened cryo-EM maps of p97^E470D^ (left) and p97^D592N^ (right) mutants, indicating the presence of ATP*γ*S density in both the D1 and D2 domains. Average sigma threshold values for p97^E470D^ and p97^D592N^ EM maps are 8.4, 8.0, respectively. **(C)** Expanded view of the D1 domain loop region (residues 425-445) of p97^WT^ (grey), p97^E470D^ (cyan) and p97^D592N^ (purple). **(D)** Schematic showing the disruption of the intra-protomer D1-D2 communication, leading to a failure of the mutant N-domain to move into the “up” conformation upon ATP*γ*S binding. **(E) Left:** p97^E470D^ protomer is displayed showing backbone RMSD from D1D2-ATP*γ*S-bound p97^WT^ [PDB:5FTN]. **Right:** p97^D592N^ protomer is displayed showing backbone RMSD from D1D2-ATP*γ*S-bound p97^WT^ [PDB:5FTN].

Although we would expect the E470D mutation to have minimal effects because of the conservative substitution, residue 470 is located at a critical juncture between the D1 and D2 domains. E470 is proximal to residues F539 in D2 and D373 in D1, and the major effect of the mutation highlights the structural role of this sidechain in maintaining the network of interactions by which the structural changes in the D2 domain are communicated to the D1 domain. D592 is located in the pore loop region of the hexamer at the inter-protomer interface (Fig. 1C), indicating that interactions distal to the D1-D2 interface can also influence the coupling between these domains. As observed during the activity assays, the p97^E470D^ and p97^D592N^ mutants differ in terms of their effects on ATPase activity (Supplementary Fig. 1A). This suggests that p97^E470D^ stimulates increased ATPase activity in the D2 domain, possibly by removing the requirement for ATP-binding to the D1 domain for D2 hydrolysis, whereas this intra-protomer D1-D2 communication across the linker is maintained in the p97^D592N^ mutant.

### p97 Inhibition and Structural Rearrangements Induced by CB-5083 Binding

CB-5083 is a potent inhibitor of p97 function, with an IC50 of ∼10 nM (24). The binding site of CB-5083 to the D2 domain was characterized in a previous X-ray crystallographic study using p97 truncated to only contain the D1 and D2 domains p97^D1D2^ structure determined at a resolution of 3.75 Å (25). In order to assess the effect of CB-5083 binding in the context of full-length protein, we carried out cryo-EM studies of p97^WT^ bound to CB-5083 in the presence of either ADP or ATPγS. The cryo-EM structure of the ADP-CB-5083-p97^WT^ complex resolved at an overall resolution 2.5 Å allows unambiguous positioning of CB-5083 in the D2 domain. The inhibitor is held in place by a combination of polar interactions mediated through its terminal amide side chain and central quinazoline scaffold, and non-polar interactions involving the pendant phenyl group. The tertiary and quaternary structures of the D1 and D2 domains remain the same under both conditions, demonstrating that when CB-5083 is bound to the D2 domain, the conformation of the D1 domain is locked and unresponsive to either the presence or the identity of the bound nucleotide. Importantly, the N-domain is in the “down” position when either ADP or ATPγS occupy the D1 nucleotide binding site (Fig.4A and Supplementary Fig.2B). Notably, the vast majority of p97 particles bound to CB-5083 were in the dodecameric state, regardless of D1 nucleotide occupancy (Supplementary Fig. 15, 16), this state is often observed only in the ADP-bound p97^WT^ structures but not in the ATP*γ*S-bound structures, further supporting the observation that the CB-5083-bound D2 domain is locked in conformation I. In this respect, the effect of CB-5083 binding is similar to that observed for the E470D and D592N mutants, where the conformation of the N and D1 domains is unchanged with binding of either ADP or ATPγS (Fig. 4B). However, unlike these mutants, CB-5083-bound p97^WT^ is locked in conformation I, and retains the same tertiary structure in the D2 domain as that seen for ADP-bound p97.

**Fig. 4:**
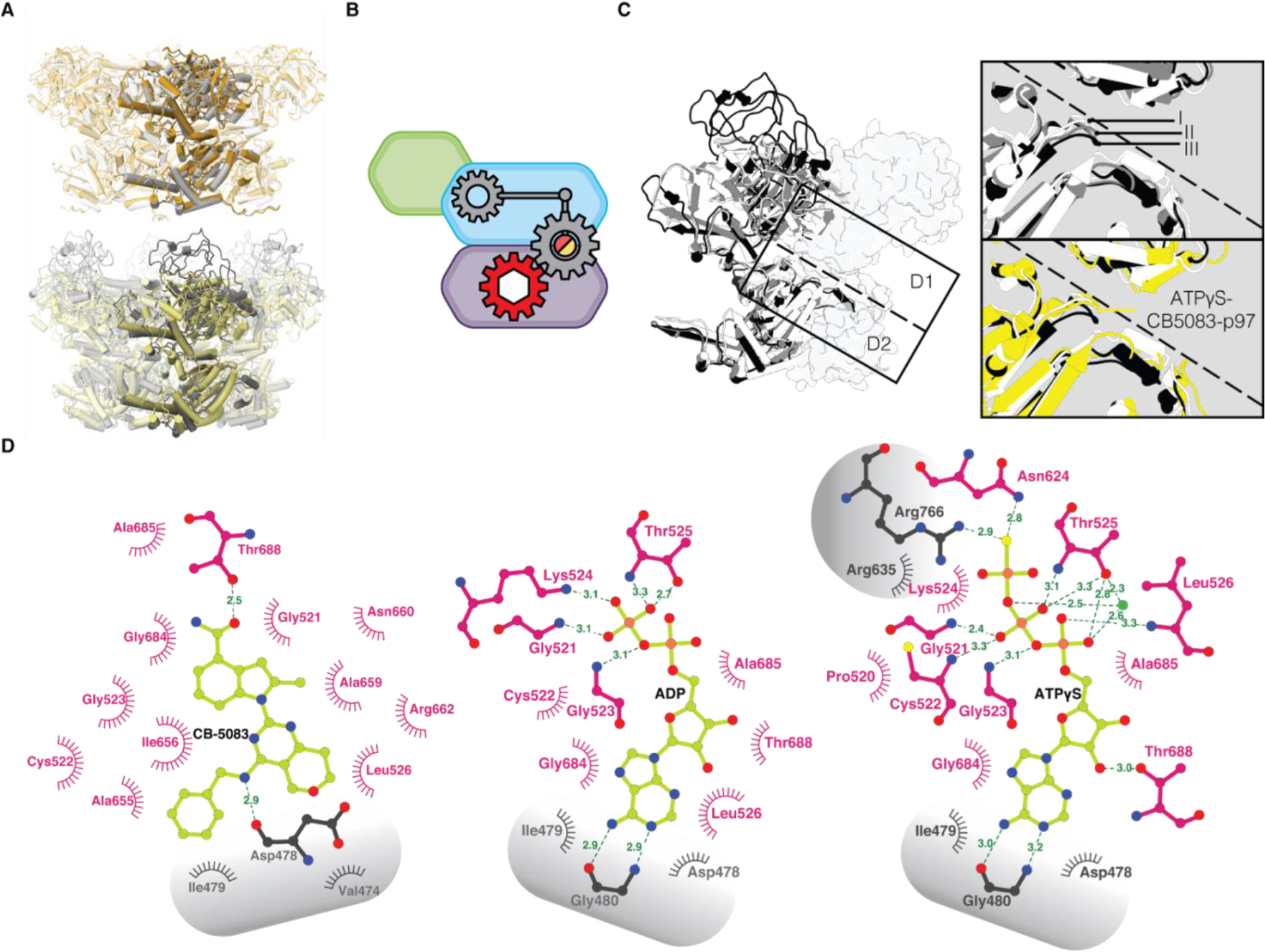
p97 Inhibition and Structural Rearrangements upon CB-5083 Binding. **(A) Top:** Superposition of ADP-bound p97^WT^ (grey) and ADP-CB-5083-bound p97^WT^ (orange) hexameric structures, with monomer highlighted. **Bottom:** Superposition of ATPγS-bound p97^WT^ (dark grey) and ATPγS-CB-5083-bound p97^WT^ (yellow) hexameric structures, with monomer highlighted. **(B)** Schematic showing the complete allosteric inhibition of p97 following CB-5083 binding independent of D1 nucleotide state, illustrating D2 allosteric activation must occur before the D1 domain can proceed to the active ATP*γ*S-bound conformation III. **(C) Left:** Overlay of conformations I, II and III of p97 monomer with adjacent monomer outlined. **Top right**: Zoom into critical D1-D2 inter-protomer interactions, illustrating the key movements in the D1 and D2 domain upon sequential ATP*γ*S-binding. **Bottom right**: Overlay of the same area with conformation II omitted for clarity and the D1-ATP*γ*S and D2-CB-5083 bound structure added in yellow, illustrating the absence of a conformational shift in the D2 domain leading to the inhibition of the allosteric activation of the D1 domain. (**D)** Ligplot of interacting residues in the binding domain from left to right of CB5083, ADP and ATP*γ*S. Interacting residues from adjacent protomers labelled in grey.

**Fig. 5:**
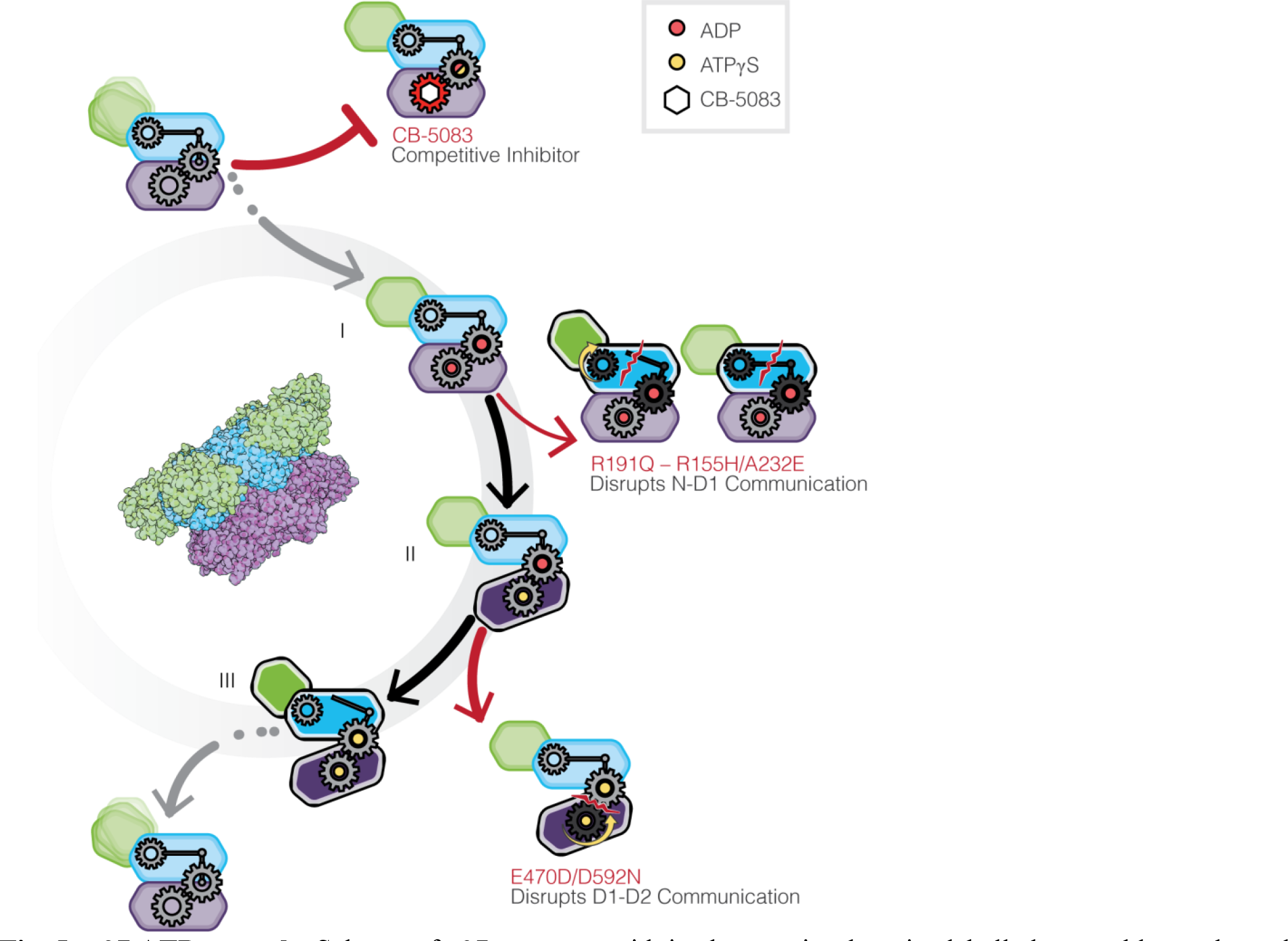
p97 ATPase cycle. Scheme of p97 protomer with its three major domains labelled green, blue and purple [N, D1 and D2 respectively], representing the four discrete stages of nucleotide binding in p97^WT^ and the effects of mutants/inhibitors on each of these stages.

CB-5083, as an ATP-competitive inhibitor, can exert its effects on p97 function by blocking the allosteric conformational changes required for transition from conformation I to conformation II (Fig. 4C). The same step is blocked by the phenyl indole based allosteric inhibitor UPCDC-30245, which binds p97 distal to the nucleotide binding site, but prevents the transition to conformation II because of a steric block of the conformational change in the D2 domain by the bound inhibitor (5).

A detailed inspection of the interactions in the nucleotide binding pocket provides an explanation for the effects observed with CB-5083 binding. As shown in Fig. 4D, the interactions of the two phosphate moieties in ADP are confined to a single nucleotide binding domain, but when ATPγS is bound, the third (gamma) phosphate moiety is involved in H-bonding interactions with R766 and R635 in the neighboring protomer. These additional interactions explain why ATPγS binding initiates the sequence of quaternary structural changes via interactions at the interprotomer interface. Although CB-5083 binding site overlaps with the nucleotide binding site of the D2 domain, and shares interactions with some of the residues that interacts with ADP/ATP, the key inter-protomer interaction between R766 and the *γ*-phosphate of ATP are missing in the CB-5083 bound structure, preventing the generation of the type of quaternary conformational changes seen upon ATPγS binding to p97^WT^.

## DISCUSSION

The cryo-EM structural studies presented here allow us to propose a structural model for the observed increase in ATPase activity of MSP-1 related p97 mutants. Our results demonstrate that the effects of the mutations are mediated chiefly through the destabilization of the N domain in the ADP-bound state, either through a loss of inter-domain interactions in p97^R155H^ and p97^R191Q^ or through inter-protomer steric effects in p97^A232E^. The increased flexibility observed in the N domain is consistent with previously published NMR studies of p97 (17). We also showed that two mutants, p97^D592N^ and p97^E470D^ act primarily through their ATP-bound states suggesting a second possible mechanism for p97 dysfunction through the disruption of interdomain communication.

It has been shown that introduction of Walker A mutants (nucleotide-binding defective) into either D1 or D2 domains (K251T or K524T, respectively) in p97^WT^ leads to an almost complete loss of ATPase activity in both domains (16). Also, introduction of Walker B mutants (ATP-hydrolysis deficient) into the D1 domain (E305Q) in p97^WT^ leads to a slight but significant increase in overall ATPase activity and a similar mutant in the D2 domains (E578Q) almost completely abolishes the activity of p97. Similar observations were made for MSP-1 mutants (12, 16), suggesting that while the ATPase activity of p97 is dependent on nucleotide binding to both D1 and D2 domains, ATP hydrolysis in the D1 domain is not necessary for p97 ATPase activity in either wild-type or mutant forms of the protein. This indicates the primary function of the D1 domain is to facilitate D2 ATPase activity through binding ATP, which leads to a conformational change in the N domain and therefore increasing the overall rate of D2 ATPase activity. As we have observed very little change in the nucleotide-binding sites in the D2 domain of the MSP-1 mutants, the previously observed increase in ADP off-rate and ATPase activity of the mutants studied may not necessarily be due to a direct conformational change in the nucleotide-binding domain (6). Instead, these changes in activity are likely because of the destabilization of the N domain in the ADP-bound conformation.

Previously, it was observed that some of these mutants destabilize the phosphate ion intermediate after ATP hydrolysis (26) in the D1 domain. It is possible that altering the conformation of the D1 domain towards an ATP-like state, decreases the half-life of the intermediate and therefore increases the concentration of ATP-bound complexes within mutant cells relative to WT, creating a quicker turnover of ATP. Our observations are in agreement with previously published NMR studies of MSP-1 mutants, whereby MSP-1 “mutations shift the ADP-bound form of the enzyme towards an ATP-like state in a manner that correlates with disease severity” (17).

The p97^E470D^ and p97^D592N^ mutations exert their effects via changes in the ATPγS-bound conformation and appear to decouple the D2 domain from the D1 domain. In the case of p97^E470D^ this decoupling leads to an increase in ATPase activity, it is possible that the requirement for ATP-binding in the D1 domain for ATP hydrolysis in the D2 domain in WT is abolished by the E470D mutation, thus allowing the D2 domain to cycle through ATP independent of D1 nucleotide state. While p97^D592N^ mutants appear to result in a similar disruption of ATP-induced conformational changes this does not translate to an effect on the rate of ATP hydrolysis. This suggests that the D2 domain ATP-hydrolysis of the D592N mutant may still rely on the communication of the nucleotide state of the D1 domain, further highlighting the importance of the D1-D2 linker (463–480) in regulating D2 ATP hydrolysis. Our results indicate that interdomain communication between D1 and D2 domains can be influenced by distal interactions communicated to the domain interface via allosteric effects. Importantly, our structural studies with CB-5083 demonstrate that although this inhibitor competes with ATP binding to the D2 domain, the net effect of inhibitor binding is to block the critical allosteric changes that occur via inter-protomer and inter-domain communication in a manner similar to that observed with inhibitors such as UPCDC-30245 (5) that do not block ADP binding to either D1 or D2 sites, but also lock p97 in conformation I as observed with CB-5083. That is, it is necessary for the D2 domain to move into the active ATP*γ*S-bound conformation II before the D1 domain can proceed to the active ATP*γ*S-bound conformation III. This conformational block upon inhibitor binding occurs in conformation I in contrast to what we observe in the CB-5083 resistant, p97^E470D^ mutant where the block occurs at conformation II, demonstrating the movement from conformation I to II is necessary to proceed to conformation III but not sufficient. Altogether, the analysis of the structures of p97 disease mutants and of the structure of p97^WT^ bound to CB-5083 provide insights into the importance of inter-protomer and inter-domain interactions that govern p97 activity (Fig. 5). The availability of atomic resolution models for disease mutants, and understanding of the similarities and differences in allosteric changes induced by mutations vs inhibitor binding provides a powerful starting point for structure-guided drug design of potent modulators whose effects can be specifically tuned for regulating p97 function.

## MATERIALS AND METHODS

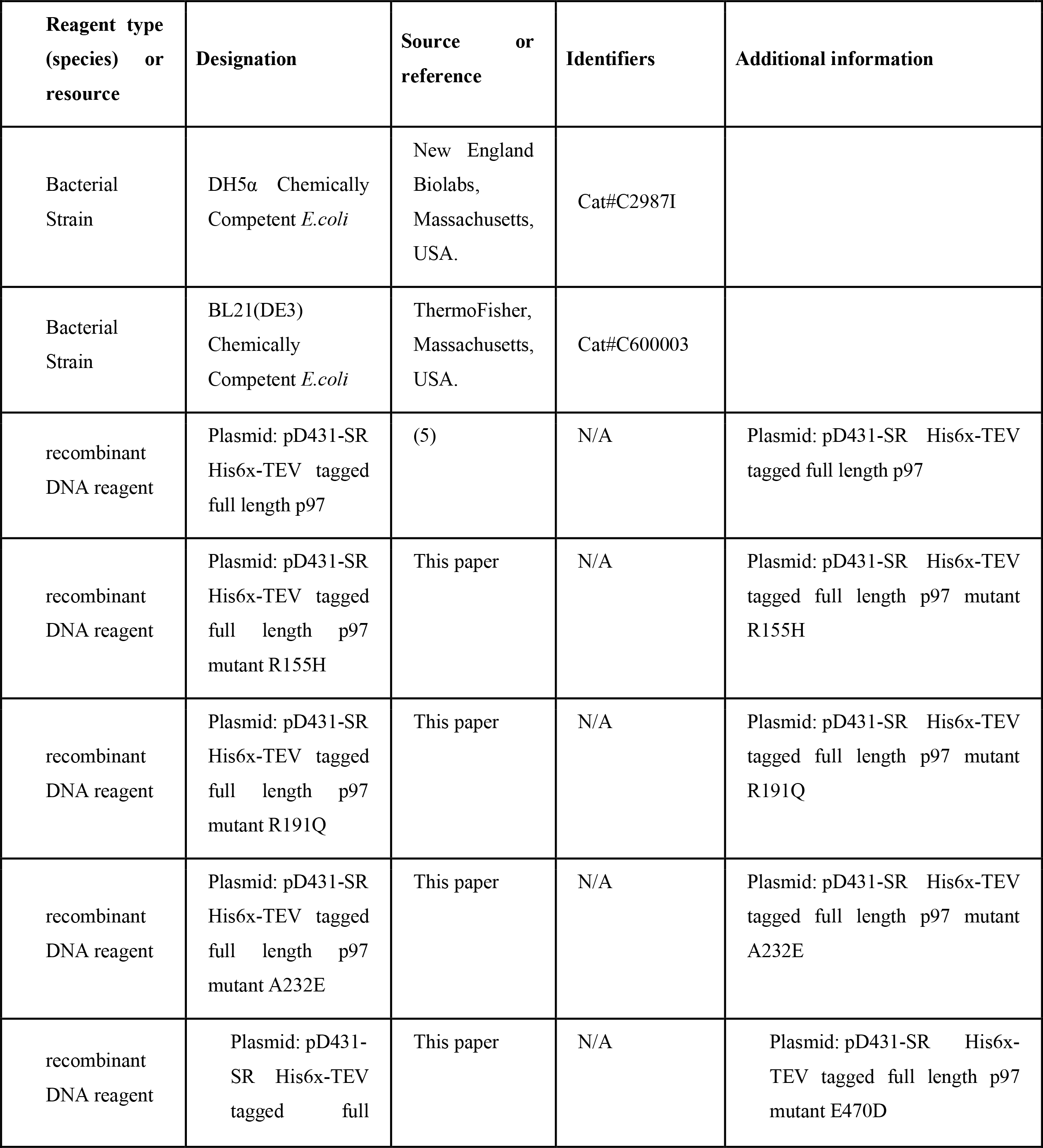
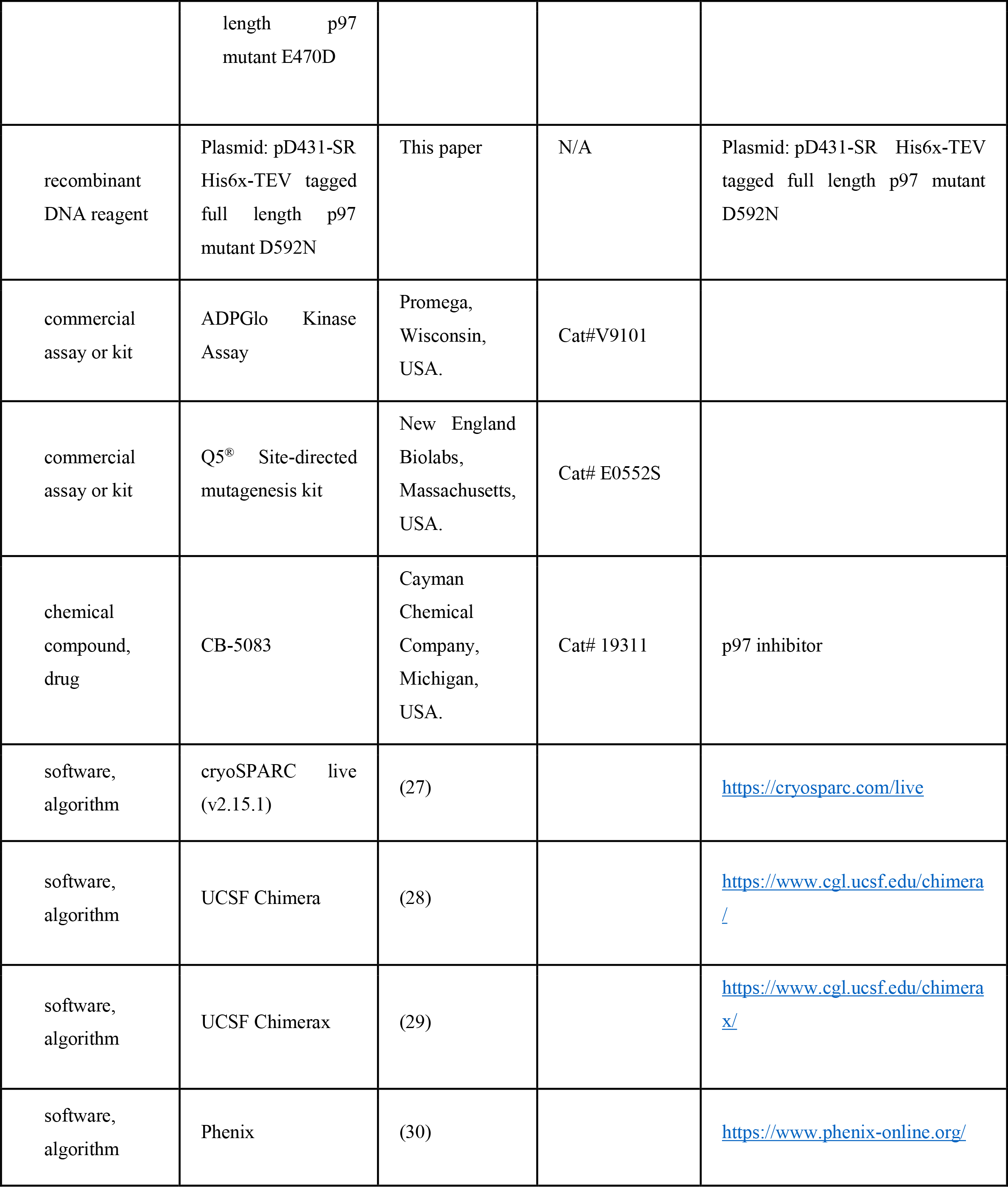

### Construction of Expression Plasmids

Multiple nucleotides were substituted at areas identified in key disease mutations: c.464G>A (R155H); c.572G>A, 573T>A (R191Q); c.695C>A (A232E); c.1410G>C (E470D); c.1774G>A (D592N) (see Table S1 for full list of primer sequences) using the Q5^®^ Site-directed mutagenesis kit (New England Biolabs, MA, USA). Each mutant sequence was verified for insertion of the correct mutation using sanger sequencing.

### Protein Expression and Purification

Full-length recombinant His6x-tagged human p97 mutants were purified with some adaptations as described in (31). Briefly, after overnight incubation at 27 °C with 0.5 mM Isopropyl β-D-1-thiogalactopyranoside (IPTG) (Fischer Scientific, MA, USA), *E. coli* BL21 (DE3) (Thermo Fischer, MA, USA) cells expressing plasmid-encoded mutant p97 were suspended in 20 ml of lysis buffer; 50 mM Tris-HCL pH 8, 300 mM NaCl, 0.5 mM β-Mercaptoethanol (BME) (Fischer Scientific, MA, USA) and 1x Protease Inhibitor Cocktail tablet (Thermo Fisher, MA, USA) and lysed. Following centrifugation, imidazole was added to the resulting supernatant to a final concentration of 20 mM and loaded onto a Ni-NTA column (Thermo Fischer, MA, USA) and eluted with 300 mM imidazole. The eluted protein was flash frozen in liquid nitrogen and stored in aliquots at -80 °C without glycerol.

### ATPase Assays

Protein concentrations were first normalized using Pierce BCA Protein Assay kit (Thermo Fischer, MA, USA), a standard series was prepared with the BSA provided (2 mg/ml - 25 μg/μl). 25 μl of each of the standard and unknown sample were added to a 96-well plate in triplicate, 200 μl of working reagent was added to each well; prepared according to manufacturer’s guidelines. The plate was covered and incubated at 37 °C for 30 mins, the plate was cooled to room temperature (RT) and the absorbance at 562 nm was measured for each well.

The ATPase activity was examined using the ADP-Glo^™^ Kinase Assay (Promega, WI, USA), as previously described in (32). The protein solution was diluted to 20 nM p97 with 20 μM ATP in triplicate in 5 μls of 1X Kinase Reaction Buffer A; 40 mM Tris pH 7.5, 20 mM MgCl2, 0.1 mg/ml BSA and incubated at 37 °C for 30 mins. After incubation, the solutions were equilibrated to RT for 5 mins and 5 μl of ADP-Glo™ Reagent was added to stop the reaction and consume the remaining ATP. The solution was incubated for 40 mins at RT. 10 μl of Kinase Detection Reagent was added and incubated for 30 mins. The sample’s luminescence was recorded using Varioskan LUX Multimode Microplate reader (Thermo Fischer, MA, USA). Each sample was averaged across the 3 measurements and plotted as mean ± standard deviation.

### Cryo-EM Grid Preparation and Data Acquisition

10 mg/ml frozen aliquots of the mutant solution were spun down at 12,000g for 10 mins and the supernatant was diluted to a concentration of 2 mg/ml in protein storage buffer with 0.5 μM octyl glucoside, 1 mM TCEP (Fischer Scientific, MA, USA) and 1 mM ADP (Sigma Aldrich, MO, USA) or 1 mM ATPγS (Sigma Aldrich, MO, USA) or 1mM CB-5083 (Cayman Chemical Company, MI, USA) depending on the experiment and incubated on ice before vitrification. Protein vitrification was achieved using the Leica GP2 plunge freezer on Quantifoil (R1.2/1.3, 200 or 300 mesh-Cu) holey carbon grids (Electron Microscopy Sciences, PA, USA). Firstly, coated grids were glow discharged at 25 mA for 15 sec and placed on a pair of tweezers in the blotting chamber. Chamber temperature was set to 20 °C with 100% humidity. 3 μl of prepared protein solution was loaded on the grid. The grid was blotted for 6 sec, followed by plunge-freezing in liquid ethane.

Frozen grids were stored in cryo grid boxes in liquid nitrogen until imaging. Digital micrographs of frozen hydrated protein particles were typically recorded at 190000 x magnification with defocus range between -1 to -2.5 μm with a total dose of 50 e^-^/Å^2^ on either a 200 kV Glacios or 300 kV Krios (ThermoFisher Scientific, MA, USA) transmission electron microscope fitted with a K3 (Gatan), Falcon3 or Falcon4 (ThermoFisher Scientific, MA, USA) direct electron detector camera. Please refer to Extended Data Table 1 for a full description of the cryo-EM data collection parameters.

### Image Processing

In general, all data processing was performed in cryoSPARC v.2.15 or v.3.0.1(27) unless stated otherwise. Motion correction in patch mode, CTF estimation in patch mode, blob particle picking (using 160 Å blob as template) and particle extraction were performed on-the-fly in cryoSPARC.

After preprocessing, particles were subjected to 2D classification and 3D heterogeneous refinement. Final 3D homogeneous refinement was done with per particle CTF estimation and aberration correction, C6 (for all the p97 mutants) or D6 (for p97 bound with CB5083 and ADP/ATPγS) symmetry applied. Resolution of the 3D map was determined according to the resulting Fourier Shell Correlation (FSC) curve at a cutoff criterion of FSC = 0.143(33). Please refer to Extended Data Table 1, for a full description and Supplementary Fig. 5-16 for graphical representation of the cryo-EM structure determination and validation analysis.

### Model Building and Structure Analysis

The p97^WT^ cryo-EM structures in the ADP (PDB ID:5FTK) or ATPγS (PDB ID: 5FTN) bound states reported previously (5) were used to build initial models for the ADP- and ATPγS-bound mutants respectively and the D1-ADP- and D2-ATP*γ*S-bound WT-p97 structure (PDB ID: 5FTM) was used as a starting model for the ATPγS-bound E470D and D592N mutants, the D1-ADP ligand was replaced with an ATPγS. Using UCSF Chimera (28), the hexamer was fit into the map to get an initial structure for real-space refinement of the atomic model in PHENIX (30). The resulting atomic model was examined using the comprehensive validation tool in PHENIX. Figures were prepared using UCSF Chimera and ChimeraX (29).

## QUANTIFICATION AND STATISTICAL ANALYSIS

**Extended Data Table 1:** Statistics of Cryo-EM Data Collection, Processing and Model Refinement.

| | p97<sup>R155H</sup>-ADP | p97<sup>R155H</sup>-ATPγS |
| --- | --- | --- |
| **Cryo-EM data collection and processing** |  |  |
| Microscope | TF Titan Krios | TF Titan Krios |
| Magnification | 105000 | 105000 |
| Voltage (kV) | 300 | 300 |
| Detector | Gatan K3 | Gatan K3 |
| Dose Rate (e<sup>-</sup>/pixel/s) | 20 | 20 |
| Total Exposure (e<sup>-</sup>/Å<sup>2</sup>) | 50 | 50 |
| Defocus Range (μm) | -1 to -2.5 | -1 to -2.5 |
| Pixel Size (Å) | 0.324 | 0.324 |
| Micrographs Collected | 4410 | 5265 |
| Symmetry Imposed | C6 | C6 |
| Particles Extracted/Final | 629745/146973 | 1293922/284092 |
| Map Sharpening B-Factor | 126 | 109.5 |
| Unmasked Resolution at 0.5/0.143 FSC (Å) | 3.9/3.2 | 2.9/2.6 |
| Masked Resolution at 0.5/0.143 FSC (Å) | 3.8/3.1 | 2.8/2.5 |
| **Model building and validation** |  |  |
| Composition (#) |  |  |
| Amino Acids | 4410 | 4422 |
| Ligands ATPγS | 0 | 12 |
| ADP | 12 | 0 |
| MG | 0 | 12 |
| Bonds (RMSD) |  |  |
| Length (Å) | 0.004 (0) | 0.004 (0) |
| Angles (°) | 1.064 (0) | 1.027 (0) |
| Mean B-Factors |  |  |
| Amino Acids | 193.88 | 146.33 |
| Ligand | 131.67 | 96.80 |
| Ramachandran |  |  |
| Favoured (%) | 94.85 | 93.84 |
| Allowed (%) | 4.88 | 6.16 |
| Outliers (%) | 0.27 | 0.00 |
| Rotamer Outliers (%) | 0.32 | 0.00 |
| Clash Score | 10.13 | 11.24 |
| C-Beta Outliers (%) | 0.00 | 0.00 |
| CaBLAM Outliers (%) | 3.44 | 3.16 |
| CC(Mask) | 0.77 | 0.82 |
| MolProbity Score | 1.88 | 1.98 |

| | p97<sup>R191Q</sup>-ADP | p97<sup>R191Q</sup>-ATPγS |
| --- | --- | --- |
| **Cryo-EM data collection and processing** |  |  |
| Microscope | TF Titan Krios | TF Titan Krios |
| Magnification | 105000 | 105000 |
| Voltage (kV) | 300 | 300 |
| Detector | Gatan K3 | Gatan K3 |
| Dose Rate (e<sup>-</sup>/pixel/s) | 20 | 20 |
| Total Exposure (e<sup>-</sup>/Å<sup>2</sup>) | 50 | 50 |
| Defocus Range (μm) | -1 to -2.5 | -1 to -2.5 |
| Pixel Size (Å) | 0.826 | 0.826 |
| Micrographs Collected | 2000 | 4797 |
| Symmetry Imposed | C6 | C6 |
| Particles Extracted/Final | 2399559/294569 | 7079943/4316075 |
| Map Sharpening B-Factor | 131.2 | 153.7 |
| Unmasked Resolution at 0.5/0.143 FSC (Å) | 3.6/3.1 | 3.1/2.8 |
| Masked Resolution at 0.5/0.143 FSC (Å) | 3.5/3.1 | 3.1/2.7 |
| **Model building and validation** |  |  |
| Composition (#) |  |  |
| Amino Acids | 4464 | 4422 |
| Ligands ATPγS | 0 | 12 |
| ADP | 12 | 0 |
| MG | 0 | 12 |
| Bonds (RMSD) |  |  |
| Length (Å) | 0.04 (0) | 0.004 (0) |
| Angles (°) | 1.045 (0) | 1.090 (12) |
| Mean B-Factors |  |  |
| Amino Acids | 170.66 | 182.41 |
| Ligand | 104.04 | 129.96 |
| Ramachandran |  |  |
| Favoured (%) | 91.60 | 92.45 |
| Allowed (%) | 8.13 | 7.55 |
| Outliers (%) | 0.27 | 0.00 |
| Rotamer Outliers (%) | 0.00 | 0.03 |
| Clash Score | 9.42 | 9.94 |
| C-Beta Outliers (%) | 0.00 | 0.00 |
| CaBLAM Outliers (%) | 5.30 | 4.94 |
| CC(Mask) | 0.81 | 0.79 |
| MolProbity Score | 2.00 | 1.99 |

| | p97<sup>A232E</sup>-ADP | p97<sup>A232E</sup>-ATPγS |
| --- | --- | --- |
| **Cryo-EM data collection and processing** |  |  |
| Microscope | TF Titan Krios | TF Titan Krios |
| Magnification | 105000 | 105000 |
| Voltage (kV) | 300 | 300 |
| Detector | Gatan K3 | Gatan K3 |
| Dose Rate (e<sup>-</sup>/pixel/s) | 20 | 20 |
| Total Exposure (e<sup>-</sup>/Å<sup>2</sup>) | 50 | 50 |
| Defocus Range (μm) | -1 to -2 | -1 to -2 |
| Pixel Size (Å) | 0.76 | 0.98 |
| Micrographs Collected | 2569 | 640 |
| Symmetry Imposed | C6 | C6 |
| Particles Extracted/Final | 74417/14388 | 310167/163041 |
| Map Sharpening B-Factor | 116 | 126.2 |
| Unmasked Resolution at 0.5/0.143 FSC (Å) | 3.3/2.7 | 3.2/2.7 |
| Masked Resolution at 0.5/0.143 FSC (Å) | 3.2/2.7 | 3.2/2.6 |
| **Model building and validation** |  |  |
| Composition (#) |  |  |
| Amino Acids | 4410 | 4422 |
| Ligands ATPγS | 0 | 12 |
| ADP | 12 | 0 |
| MG | 0 | 12 |
| Bonds (RMSD) |  |  |
| Length (Å) | 0.005 (0) | 0.004 (0) |
| Angles (°) | 1.080 (12) | 1.025 (0) |
| Mean B-Factors |  |  |
| Amino Acids | 153.36 | 189.3 |
| Ligand | 97.94 | 132.17 |
| Ramachandran |  |  |
| Favoured (%) | 95.96 | 94.41 |
| Allowed (%) | 3.76 | 5.32 |
| Outliers (%) | 0.27 | 0.27 |
| Rotamer Outliers (%) | 0.03 | 0.00 |
| Clash Score | 10.02 | 7.42 |
| C-Beta Outliers (%) | 0.00 | 0.00 |
| CaBLAM Outliers (%) | 3.44 | 4.39 |
| CC(Mask) | 0.80 | 0.79 |
| MolProbity Score | 1.8 | 1.79 |

| | p97<sup>E470D</sup>-ADP | p97<sup>E470D</sup>-ATPγS |
| --- | --- | --- |
| **Cryo-EM data collection and processing** |  |  |
| Microscope | TF Titan Krios | TF Titan Krios |
| Magnification | 105000 | 105000 |
| Voltage (kV) | 300 | 300 |
| Detector | Gatan K3 | Gatan K3 |
| Dose Rate (e<sup>-</sup>/pixel/s) | 20 | 20 |
| Total Exposure (e<sup>-</sup>/Å<sup>2</sup>) | 50 | 50 |
| Defocus Range (μm) | -1 to -2.5 | -1 to -2.5 |
| Pixel Size (Å) | 0.415 | 0.415 |
| Micrographs Collected | 4653 | 4810 |
| Symmetry Imposed | C6 | C6 |
| Particles Extracted/Final | 1180899/283837 | 1329693/343347 |
| Map Sharpening B-Factor | 106.2 | 94.2 |
| Unmasked Resolution at 0.5/0.143 FSC (Å) | 3.5/2.9 | 3.0/2.5 |
| Masked Resolution at 0.5/0.143 FSC (Å) | 3.4/2.7 | 2.9/2.5 |
| **Model building and validation** |  |  |
| Composition (#) |  |  |
| Amino Acids | 4410 | 4368 |
| Ligands ATPγS | 0 | 12 |
| ADP | 12 | 0 |
| MG | 0 | 12 |
| Bonds (RMSD) |  |  |
| Length (Å) | 0.002 (0) | 0.003 (0) |
| Angles (°) | 0.672 (42) | 0.753 (12) |
| Mean B-Factors |  |  |
| Amino Acids | 147.89 | 128.29 |
| Ligand | 111.01 | 81.02 |
| Ramachandran |  |  |
| Favoured (%) | 96.03 | 94.31 |
| Allowed (%) | 3.83 | 5.41 |
| Outliers (%) | 0.14 | 0.28 |
| Rotamer Outliers (%) | 0.00 | 0.08 |
| Clash Score | 7.65 | 13.09 |
| C-Beta Outliers (%) | 0.00 | 0.00 |
| CaBLAM Outliers (%) | 3.58 | 4.58 |
| CC(Mask) | 0.78 | 0.80 |
| MolProbity Score | 1.69 | 2.01 |

| | p97<sup>D592N</sup>-ADP | p97<sup>D592N</sup>-ATPγS |
| --- | --- | --- |
| **Cryo-EM data collection and processing** |  |  |
| Microscope | TF Titan Krios | TF Titan Krios |
| Magnification | 105000 | 150000 |
| Voltage (kV) | 300 | 300 |
| Detector | Gatan K3 | Falcon 4 |
| Dose Rate (e<sup>-</sup>/pixel/s) | 20 | 6 |
| Total Exposure (e<sup>-</sup>/Å<sup>2</sup>) | 50 | 40 |
| Defocus Range (μm) | -1 to -2.5 | -1 to -2.5 |
| Pixel Size (Å) | 0.324 | 0.5 |
| Micrographs Collected | 6940 | 4464 |
| Symmetry Imposed | C6 | C6 |
| Particles Extracted/Final | 697655/118863 | 299730/175007 |
| Map Sharpening B-Factor | 104.3 | 77.1 |
| Unmasked Resolution at 0.5/0.143 FSC (Å) | 3.7/3.0 | 3.2/2.3 |
| Masked Resolution at 0.5/0.143 FSC (Å) | 3.6/2.9 | 3.0/2.2 |
| **Model building and validation** |  |  |
| Composition (#) |  |  |
| Amino Acids | 4410 | 4368 |
| Ligands ATPγS | 0 | 12 |
| ADP | 12 | 0 |
| MG | 0 | 12 |
| Bonds (RMSD) |  |  |
| Length (Å) | 0.003 (6) | 0.003 (0) |
| Angles (°) | 0.744 (12) | 0.785 (24) |
| Mean B-Factors |  |  |
| Amino Acids | 172.28 | 129.50 |
| Ligand | 141.24 | 80.95 |
| Ramachandran |  |  |
| Favoured (%) | 95.37 | 93.51 |
| Allowed (%) | 4.63 | 6.22 |
| Outliers (%) | 0.00 | 0.28 |
| Rotamer Outliers (%) | 0.00 | 0.16 |
| Clash Score | 10.54 | 12.89 |
| C-Beta Outliers (%) | 0.00 | 0.00 |
| CaBLAM Outliers (%) | 3.99 | 5.00 |
| CC(Mask) | 0.79 | 0.79 |
| MolProbity Score | 1.86 | 2.05 |

| | p97-CB-5083-ADP | p97-CB-5083-ATPγS |
| --- | --- | --- |
| **Cryo-EM data collection and processing** |  |  |
| Microscope | TF Titan Krios | TF Titan Krios |
| Magnification | 150000 | 150000 |
| Voltage (kV) | 300 | 300 |
| Detector | Falcon 4 | Falcon 4 |
| Dose Rate (e<sup>-</sup>/pixel/s) | 6 | 6 |
| Total Exposure (e<sup>-</sup>/Å<sup>2</sup>) | 40 | 40 |
| Defocus Range (μm) | -1 to -2.5 | -1 to -2.5 |
| Pixel Size (Å) | 0.5 | 0.5 |
| Micrographs Collected | 3600 | 12549 |
| Symmetry Imposed | D6 | D6 |
| Particles Extracted/Final | 204310/85876 | 727939/126510 |
| Map Sharpening B-Factor | 116.8 | 148.1 |
| Unmasked Resolution at 0.5/0.143 FSC (Å) | 3.2/2.6 | 4.2/3.5 |
| Masked Resolution at 0.5/0.143 FSC (Å) | 3.0/2.4 | 4.2/3.4 |
| **Model building and validation** |  |  |
| Composition (#) |  |  |
| Amino Acids | 8820 | 8736 |
| Ligands ATPγS | 0 | 12 |
| ADP | 12 | 0 |
| MG | 0 | 24 |
| CB-5083 (JDP) | 12 | 12 |
| Bonds (RMSD) |  |  |
| Length (Å) | 0.004 (0) | 0.004 (0) |
| Angles (°) | 0.826 (36) | 1.107 (74) |
| Mean B-Factors |  |  |
| Amino Acids | 136.46 | 216.93 |
| Ligand | 101.41 | 117.48 |
| Ramachandran |  |  |
| Favoured (%) | 94.40 | 93.22 |
| Allowed (%) | 5.32 | 6.23 |
| Outliers (%) | 0.27 | 0.55 |
| Rotamer Outliers (%) | 0.00 | 0.00 |
| Clash Score | 17.43 | 15.29 |
| C-Beta Outliers (%) | 0.00 | 0.15 |
| CaBLAM Outliers (%) | 3.71 | 5.28 |
| CC(Mask) | 0.82 | 0.79 |
| MolProbity Score | 2.12 | 2.13 |

## ACKNOWLEDGEMENTS

We thank members of the Subramaniam laboratory for helpful discussions. We thank Oda Schiøtz for help with grid preparation and vitrification in the early phase of the project. A portion of this research was supported by NIH grant U24GM129547 and performed at the PNCC at OHSU and accessed through EMSL (grid.436923.9), a DOE Office of Science User Facility sponsored by the Office of Biological and Environmental Research. The density maps and refined atomic models have been deposited in the Electron Microscopy Data Bank with accession numbers EMD-24518, 24519, 24522, 24523, 24524, 24525, 24526, 24528, 24529, 24530, 24531 and 24532 and in the Protein Data Bank with matching accession numbers of PDB-7RL6, 7RL7, 7RL9, 7RLA, 7RLB, 7RLC, 7RLD, 7RLF, 7RLG, 7RLH, 7RLI and 7RLJ respectively, for R155H-ADP, R155H-ATP*γ*S, R191Q-ADP, R191Q-ATP*γ*S, A232E-ADP, A232E-ATP*γ*S, E470D-ADP, E470D-ATP*γ*S, D592N-ADP, D592N-ATP*γ*S mutant p97, and for p97 bound to CB-5083-ADP and CB-5083-ATP*γ*S.

## SUPPLEMENTAL FIGURES

**Fig. S1:**
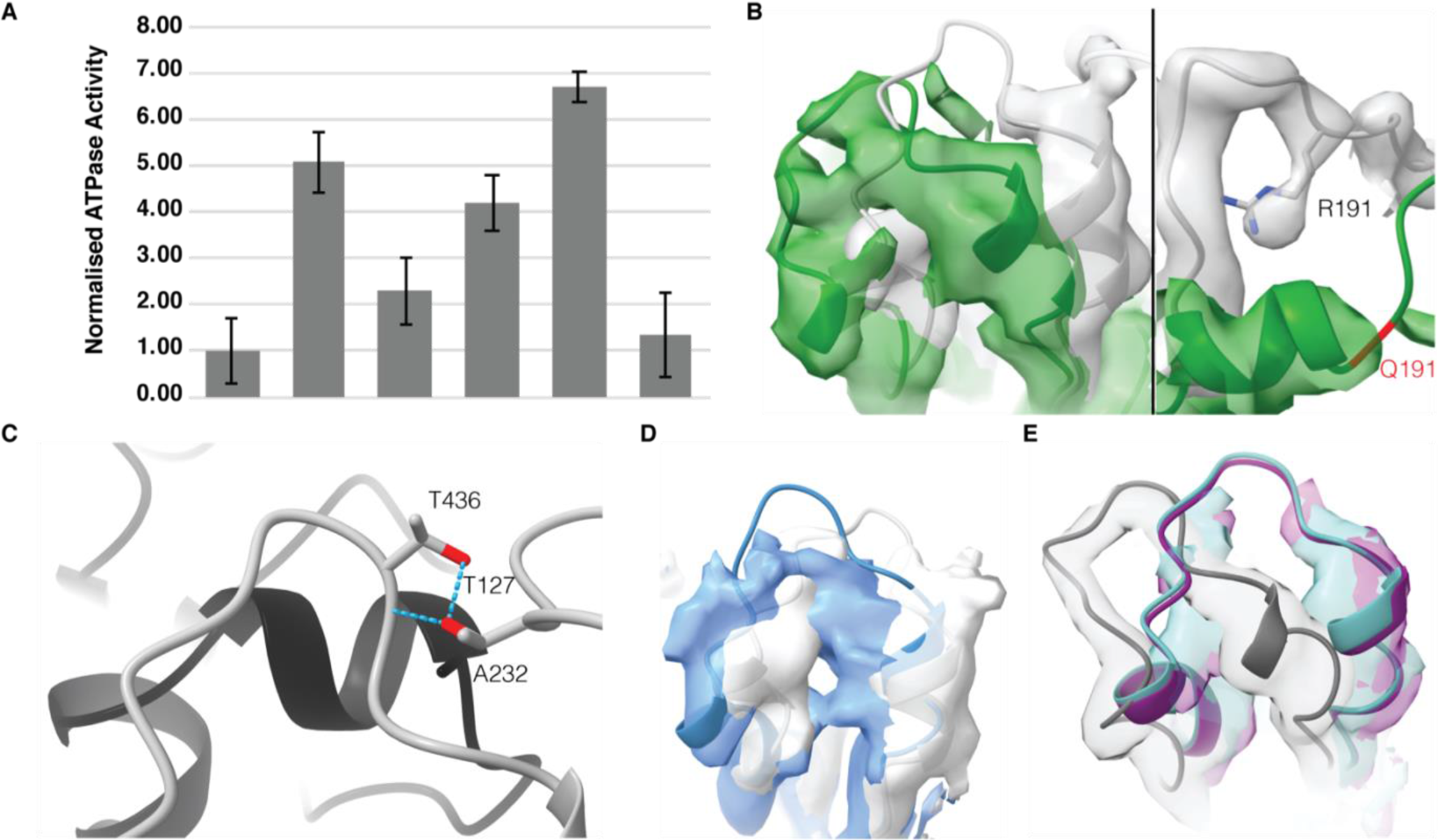
p97 Disease Mutants. **(A)** ATPase activity assay of BCA normalized concentrations of wild-type p97 and its mutants (n = 3), indicating each mutant has elevated ATPase activity relative to WT protein which is comparable with previously published data (Niwa et al., 2012). Sample size (n) represents number of averaged independent measurements in triplicate, and values are shown as mean ± standard deviation. **(B)** Overlaid ADP-bound densities of p97^WT^ (grey) and p97^R191Q^ (green) for fig.2 B+E, indicating the shift in mutant density relative to WT, sigma threshold values for mutant and WT in the left panel are 3.7 and 3.8 respectively and in the right panel 3.3 and 3.4. **(C)** Side view of A232 interacting residues between N and D1 domains in WT [PDB:5FTK], adjacent protomers labelled grey and black, tentative hydrogen bonds modelled in chimera, according to precise geometric constraints based on (Mills, 1996) labelled in cyan. **(D)** Overlaid ADP-bound densities of p97^WT^ (grey) and p97^A232E^ (green), indicating a similar shift to p97^R191Q^ in the loop region between K425 and L445 of the D1 domain in p97^A232E^ mutant density relative to WT, sigma threshold values for mutant and WT are 3.2 and 3.2 respectively. **(E)** Overlaid ATP*γ*S-bound densities of p97^WT^ (grey), p97^E470D^ (cyan) and p97^D592N^ (purple), indicating a shift in the loop region between K425 and L445 of the D1 domain in mutant density relative to WT, sigma threshold values for p97^E470D^, p97^D592N^ mutant and WT are 4.2, 4.2 and 4.2 respectively.

**Fig. S2:**
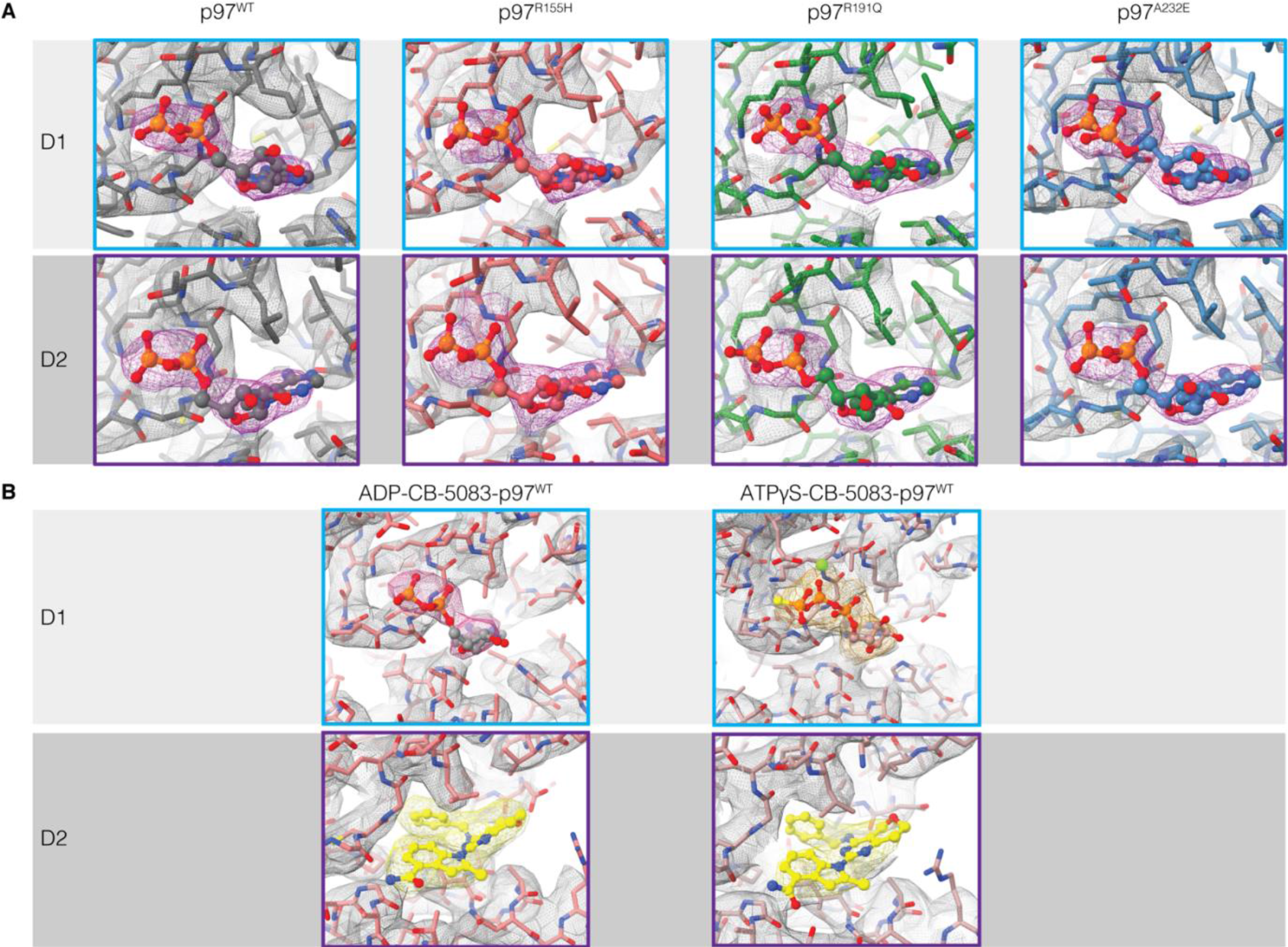
Nucleotide density in the D1 and D2 domains of the unsharpened cryo-EM maps of p97^WT^ and mutants. **(A)** Expanded view of nucleotide density in Fig. 2 models of p97^WT^ (grey), p97^R155H^ (red), p97^R191Q^ (green) and p97^A232E^ (blue) mutants, indicating the presence of ADP density in both the D1 and D2 domains. Average sigma threshold values for EM maps from left to right are 5.6, 2.9, 8.1 and 8.0. **(B)** Expanded view of nucleotide density in Fig. 4 models of p97^WT^ in the D1 and D2 domains of the unsharpened cryo-EM maps of CB5083-bound p97 with either ADP or ATP*γ*S nucleotide. Average sigma threshold values for ADP-CB-5083-bound and ATP*γ*S-CB5083-bound EM maps are 6.4 and 4.6 respectively.

**Fig. S3:**
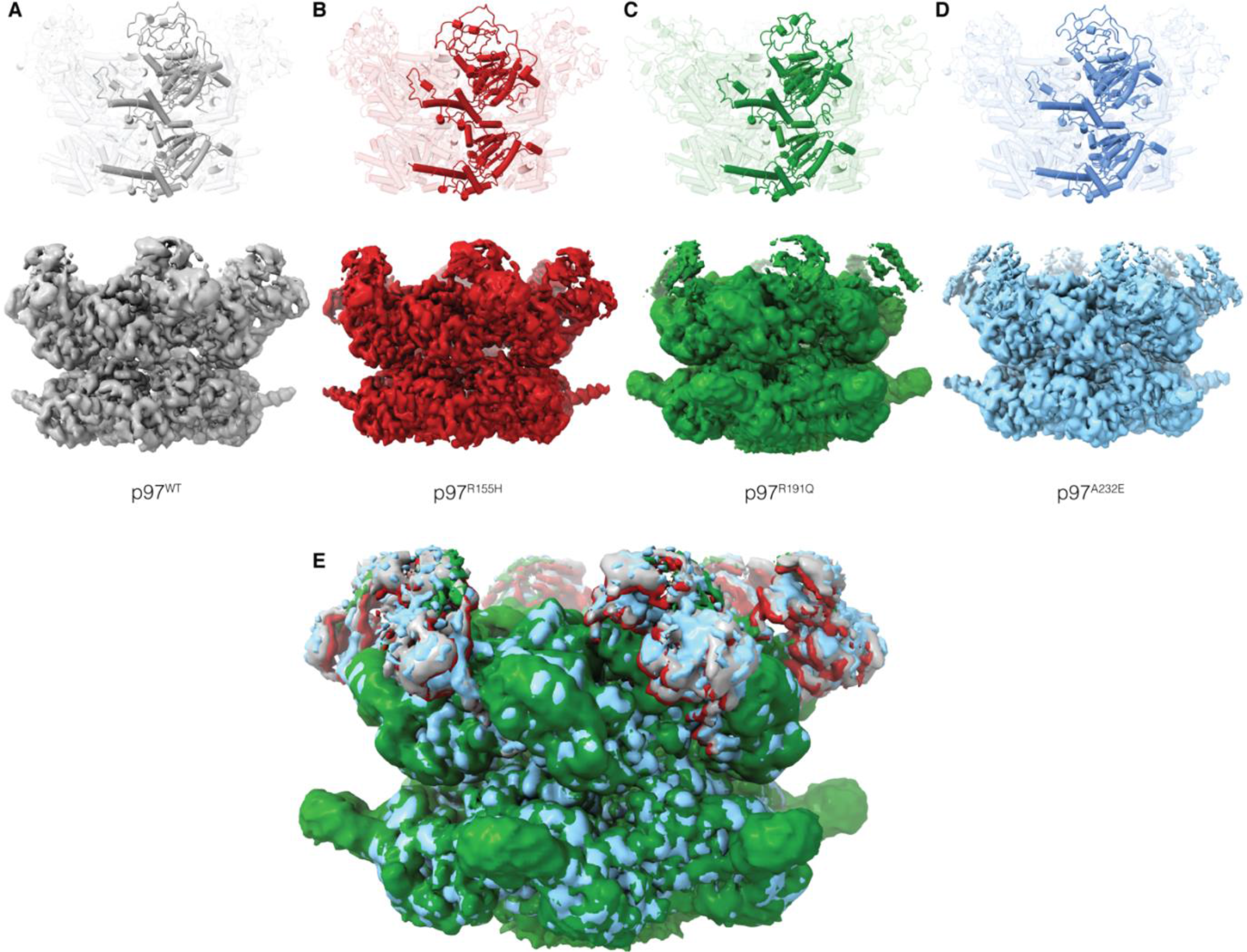
Cryo-EM Structures and their corresponding maps of ATP*γ*S-Bound p97 and Disease Mutants R155H, R191Q and A232E. ATP*γ*S-bound p97 structures, indicate similar quarternary states, with the N-domain in the “Up” position. This suggests the mutants, **(B)** p97^R155H^, **(C)** p97^R191Q^ and **(D)** p97^A232E^ have little effect on the ATP*γ*S-bound quarternary structure of p97. **(E)** Overlay of EM densities highlighting similarity in structures. Sigma threshold values for structures are 3.2, 2.6, 0.7 and 2.2 respectively.

**Fig. S4:**
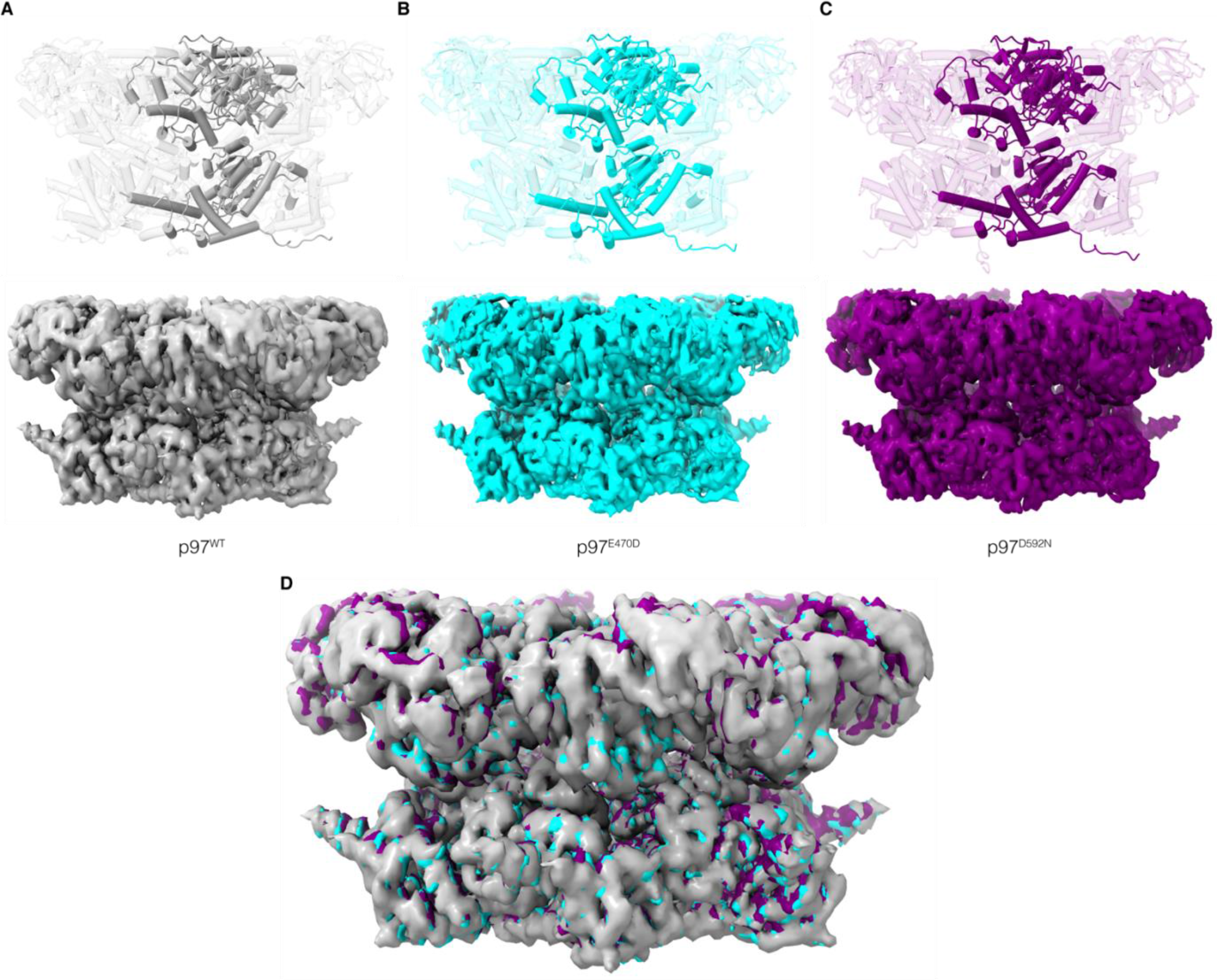
Cryo-EM Structures and their corresponding maps of ADP-Bound p97 and the Disease Mutants p97^E470D^ and p97^D592N^. ADP-bound p97 structures, indicate similar quarternary states, with the N-domain in the “Down” position. This suggests the mutants **(B)** p97^E470D^ and **(C)** p97^D592N^ have little effect on the ADP-bound quarternary structure of p97. **(D)** Overlay of EM densities highlighting similarity in structures. Sigma threshold values for structures are 2.7, 4.3, and 1.8 respectively.

**Fig. S5:**
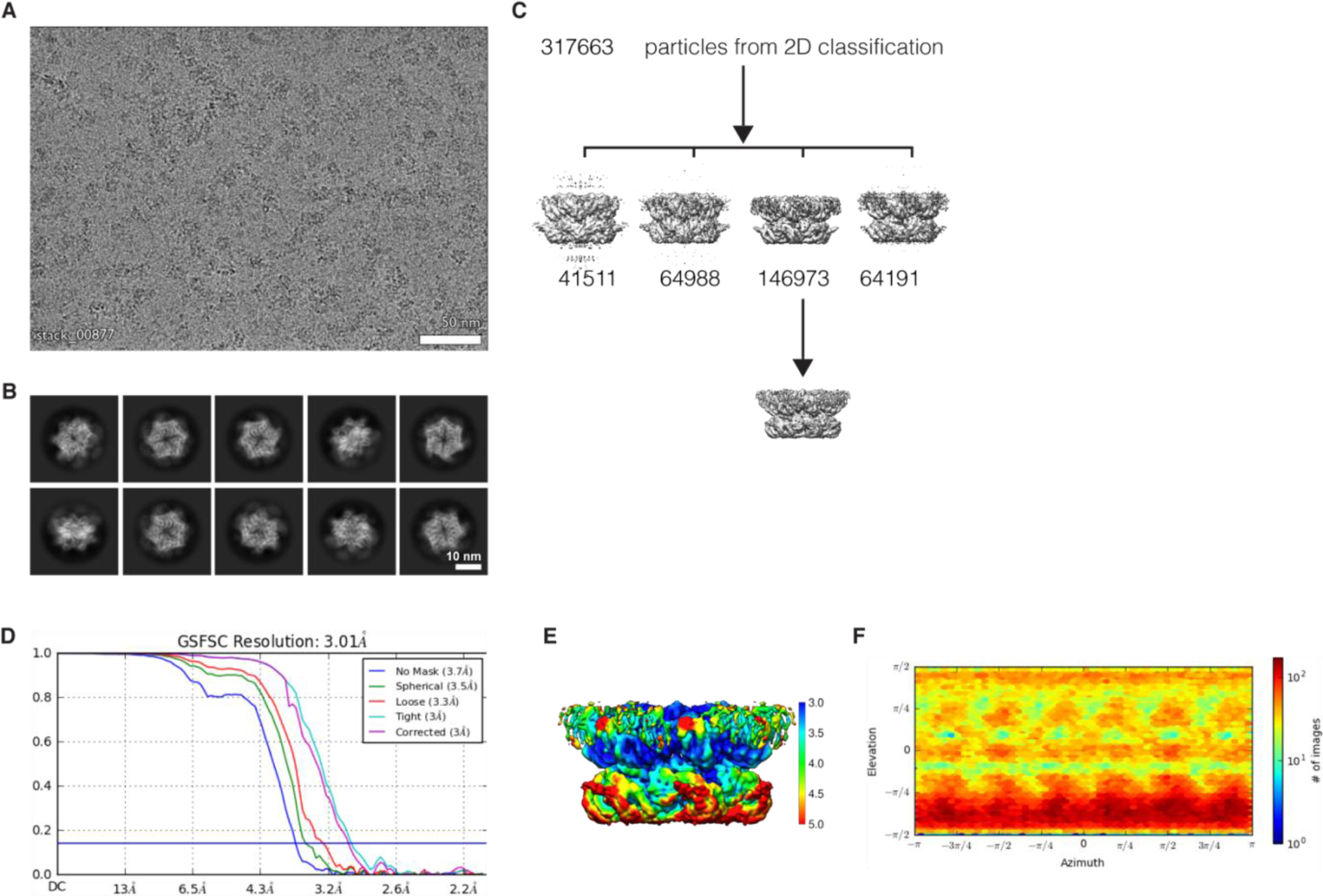
ADP-bound p97^R155H^. **(A)** A representative cryo-EM micrograph of ADP-bound p97^R155H^. **(B)** Representative 2D classes, showing multiple orientations. **(C)** Workflow of cryo-EM image processing. **(D)** FSC curves. **(E)** Cryo-EM side-view density of p97 mutants, colored according to local resolution and **(F)** Viewing direction distribution plot for mutant density.

**Fig. S6:**
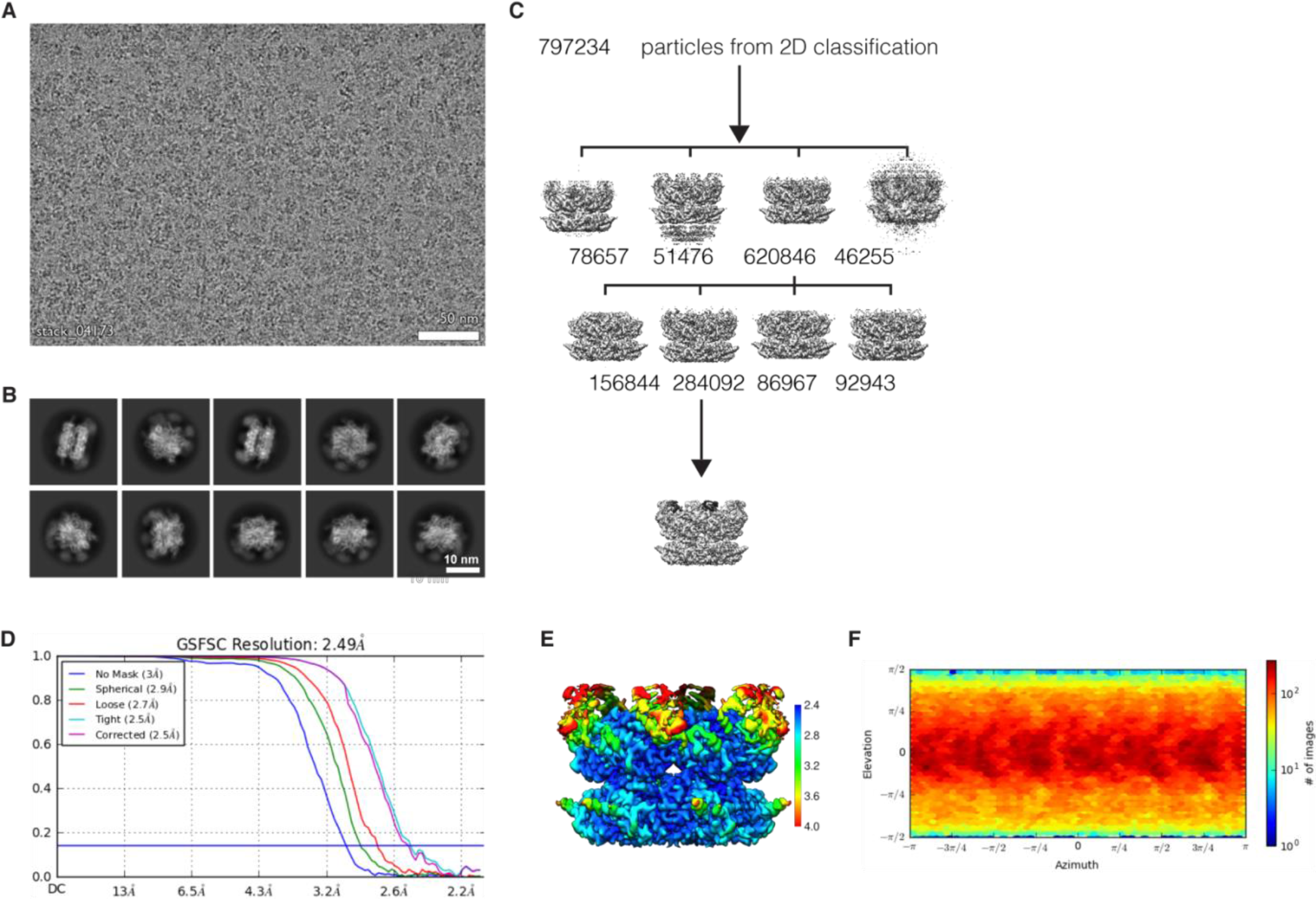
ATP*γ*S-bound p97^R155H^. **(A)** A representative cryo-EM micrograph of ATP*γ*S-bound p97^R155H^. **(B)** Representative 2D classes, showing multiple orientations. **(C)** Workflow of cryo-EM image processing. **(D)** FSC curves. **(E)** Cryo-EM side-view density of p97 mutants, colored according to local resolution and **(F)** Viewing direction distribution plot for mutant density.

**Fig. S7:**
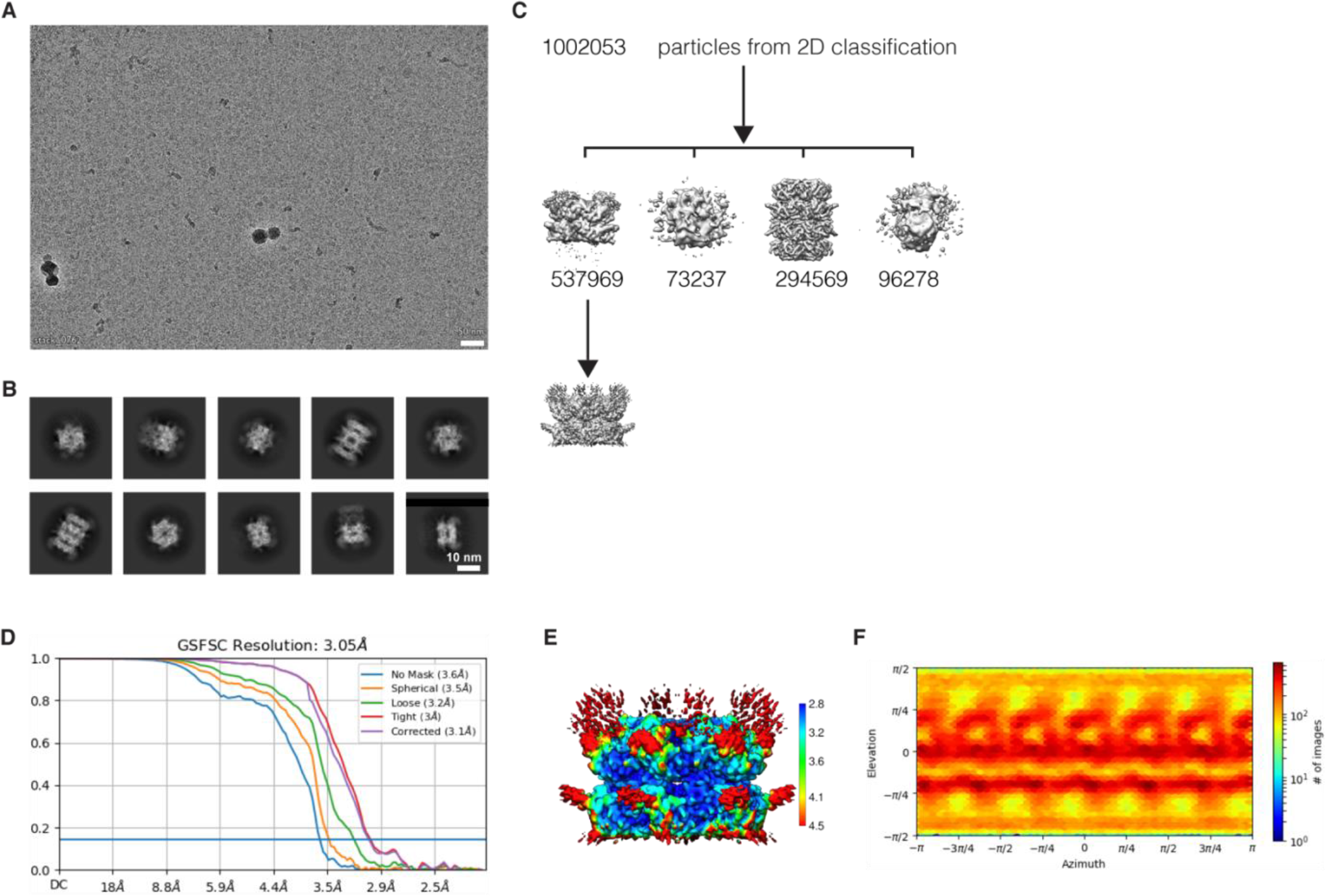
ADP-bound p97^R191Q^. **(A)** A representative cryo-EM micrograph of ADP-bound p97^R191Q^. **(B)** Representative 2D classes, showing multiple orientations. **(C)** Workflow of cryo-EM image processing. **(D)** FSC curves. **(E)** Cryo-EM side-view density of p97 mutants, colored according to local resolution and **(F)** Viewing direction distribution plot for mutant density.

**Fig. S8:**
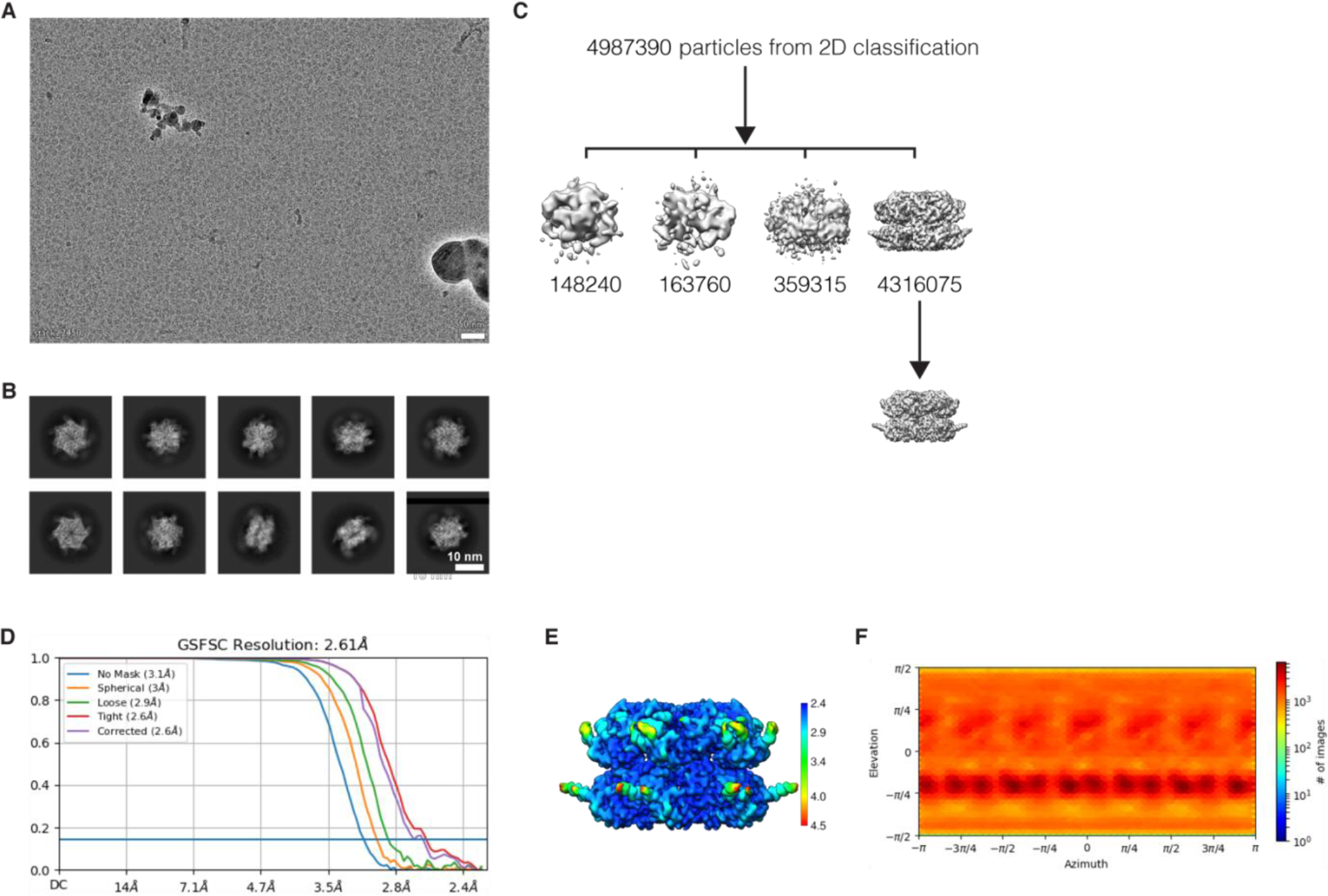
ATP*γ*S-bound p97^R191Q^. **(A)** A representative cryo-EM micrograph of ATP*γ*S-bound p97^R191Q^. **(B)** Representative 2D classes, showing multiple orientations. **(C)** Workflow of cryo-EM image processing. **(D)** FSC curves. **(E)** Cryo-EM side-view density of p97 mutants, colored according to local resolution and **(F)** Viewing direction distribution plot for mutant density.

**Fig. S9:**
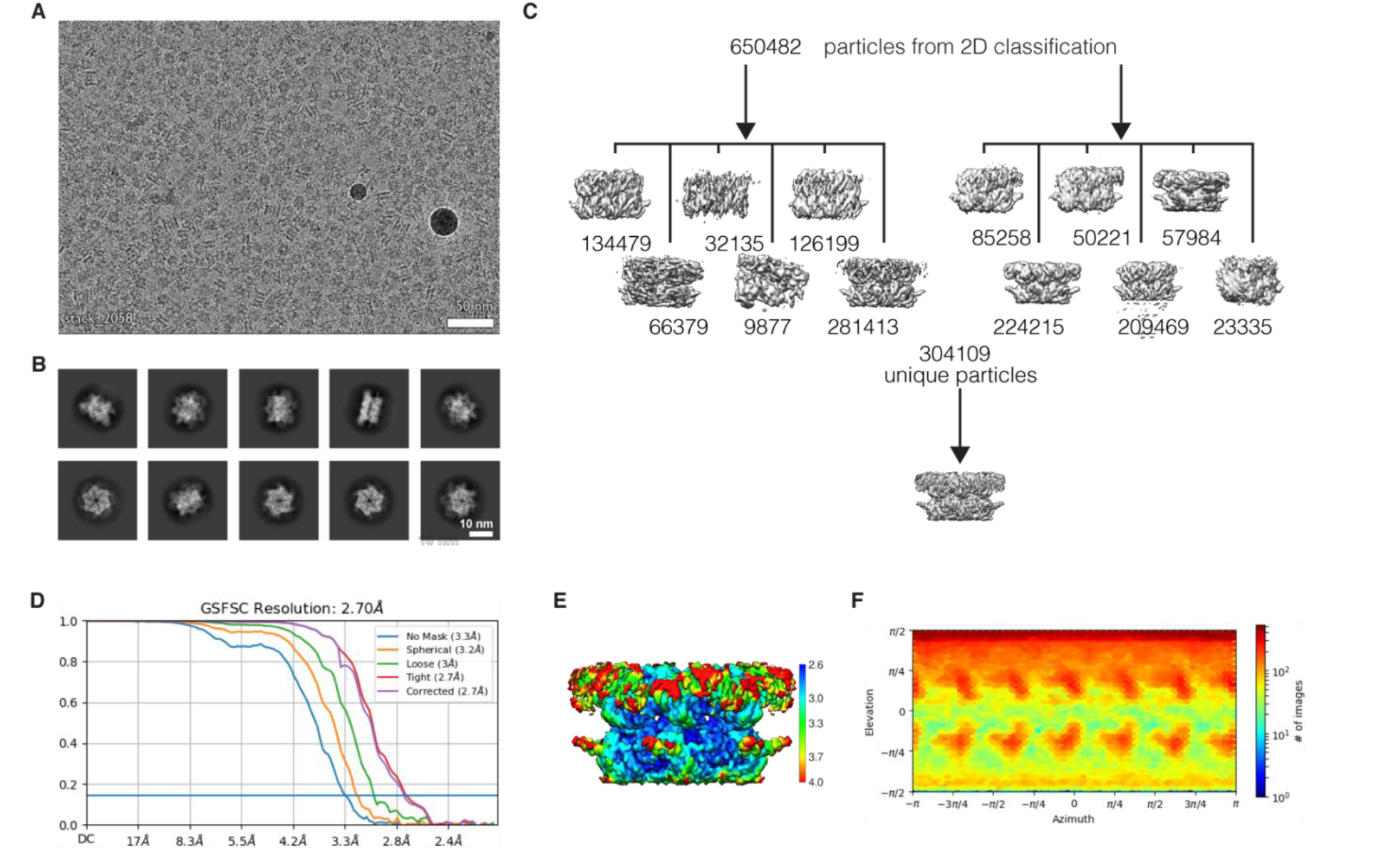
ADP-bound p97^A232E^. **(A)** A representative cryo-EM micrograph of ADP-bound p97^A232E^. **(B)** Representative 2D classes, showing multiple orientations. **(C)** Workflow of cryo-EM image processing. **(D)** FSC curves. **(E)** Cryo-EM side-view density of p97 mutants, colored according to local resolution and **(F)** Viewing direction distribution plot for mutant density.

**Fig. S10:**
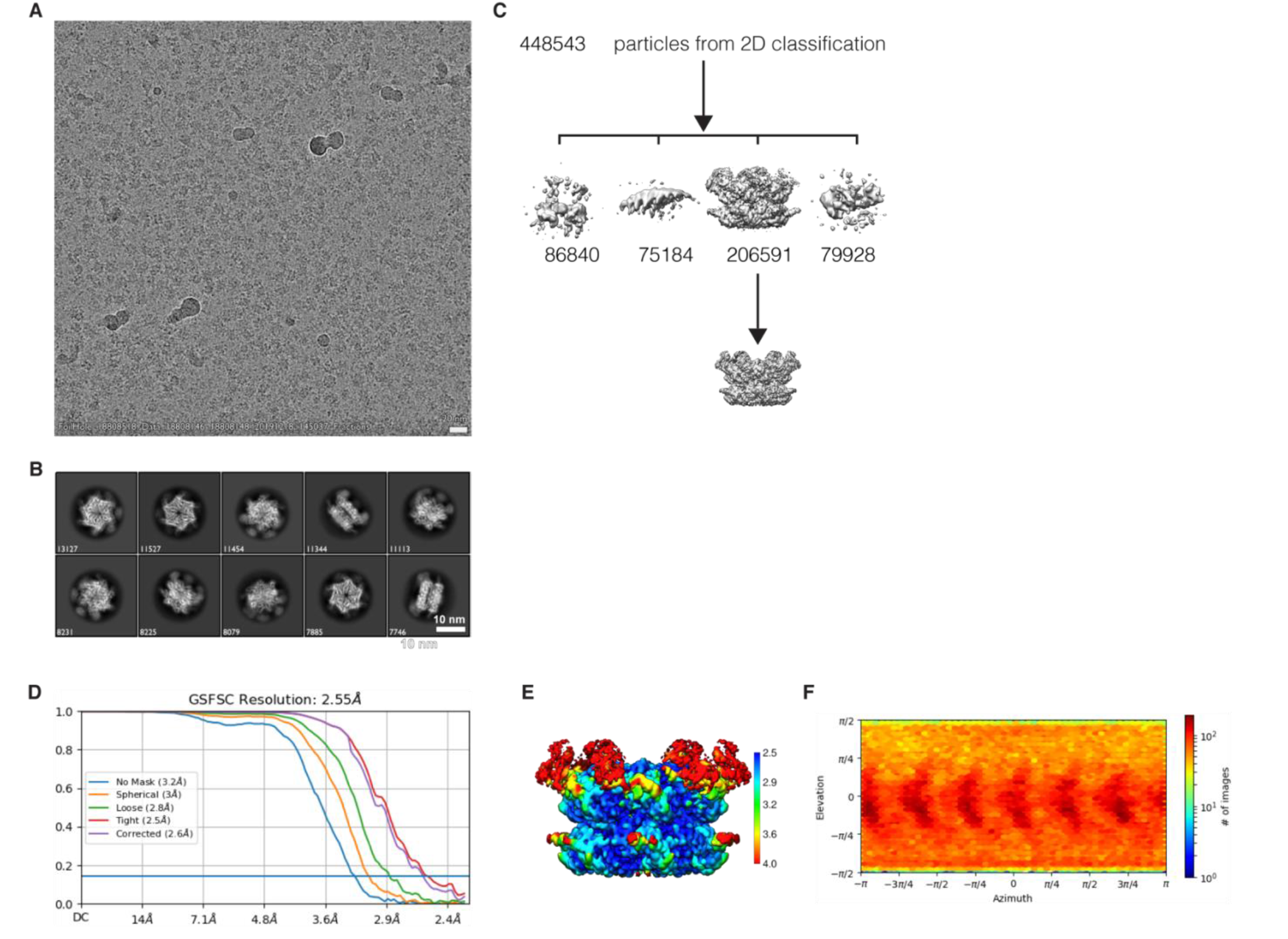
ATP*γ*S-bound p97^A232E^. **(A)** A representative cryo-EM micrograph of ATP*γ*S-bound p97^A232E^. **(B)** Representative 2D classes, showing multiple orientations. **(C)** Workflow of cryo-EM image processing. **(D)** FSC curves. **(E)** Cryo-EM side-view density of p97 mutants, colored according to local resolution and **(F)** Viewing direction distribution plot for mutant density.

**Fig. S11:**
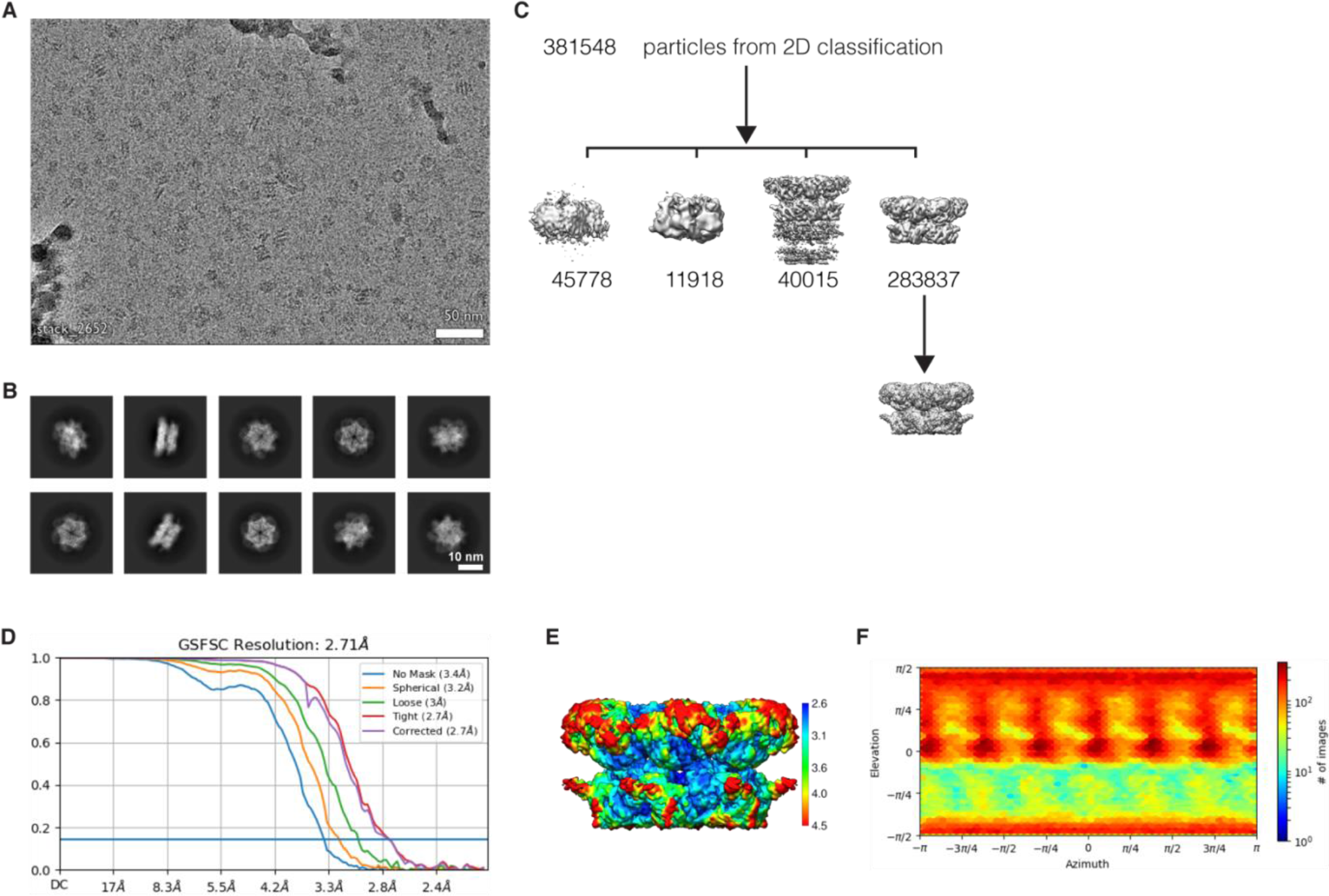
ADP-bound p97^E470D^. **(A)** A representative cryo-EM micrograph of ADP-bound p97^E470D^. **(B)** Representative 2D classes, showing multiple orientations. **(C)** Workflow of cryo-EM image processing. **(D)** FSC curves. **(E)** Cryo-EM side-view density of p97 mutants, colored according to local resolution and **(F)** Viewing direction distribution plot for mutant density.

**Fig. S12:**
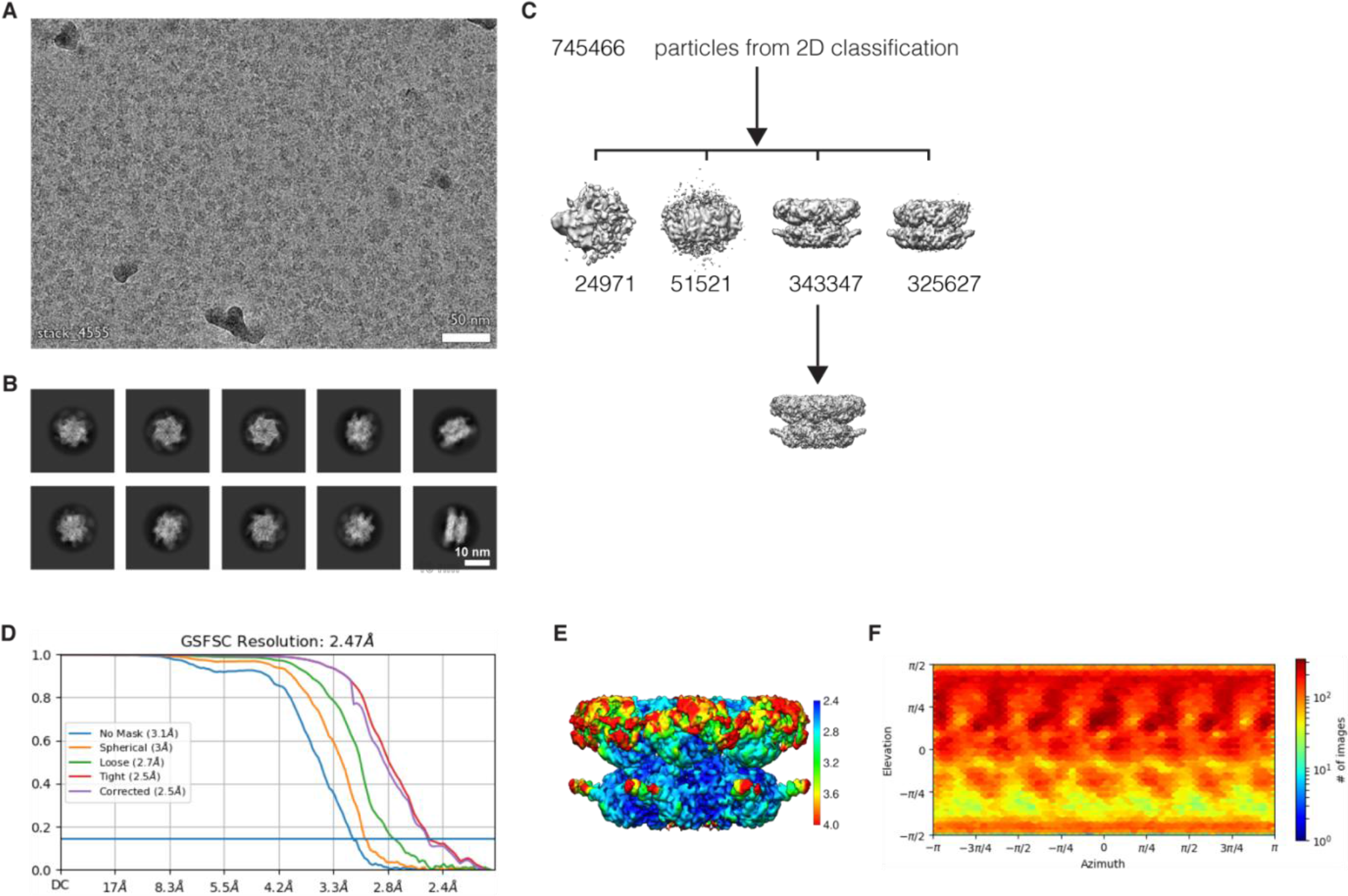
ATP*γ*S-bound p97^E470D^. **(A)** A representative cryo-EM micrograph of ATP*γ*S-bound p97^E470D^. **(B)** Representative 2D classes, showing multiple orientations. **(C)** Workflow of cryo-EM image processing. **(D)** FSC curves. **(E)** Cryo-EM side-view density of p97 mutants, colored according to local resolution and **(F)** Viewing direction distribution plot for mutant density.

**Fig. S13:**
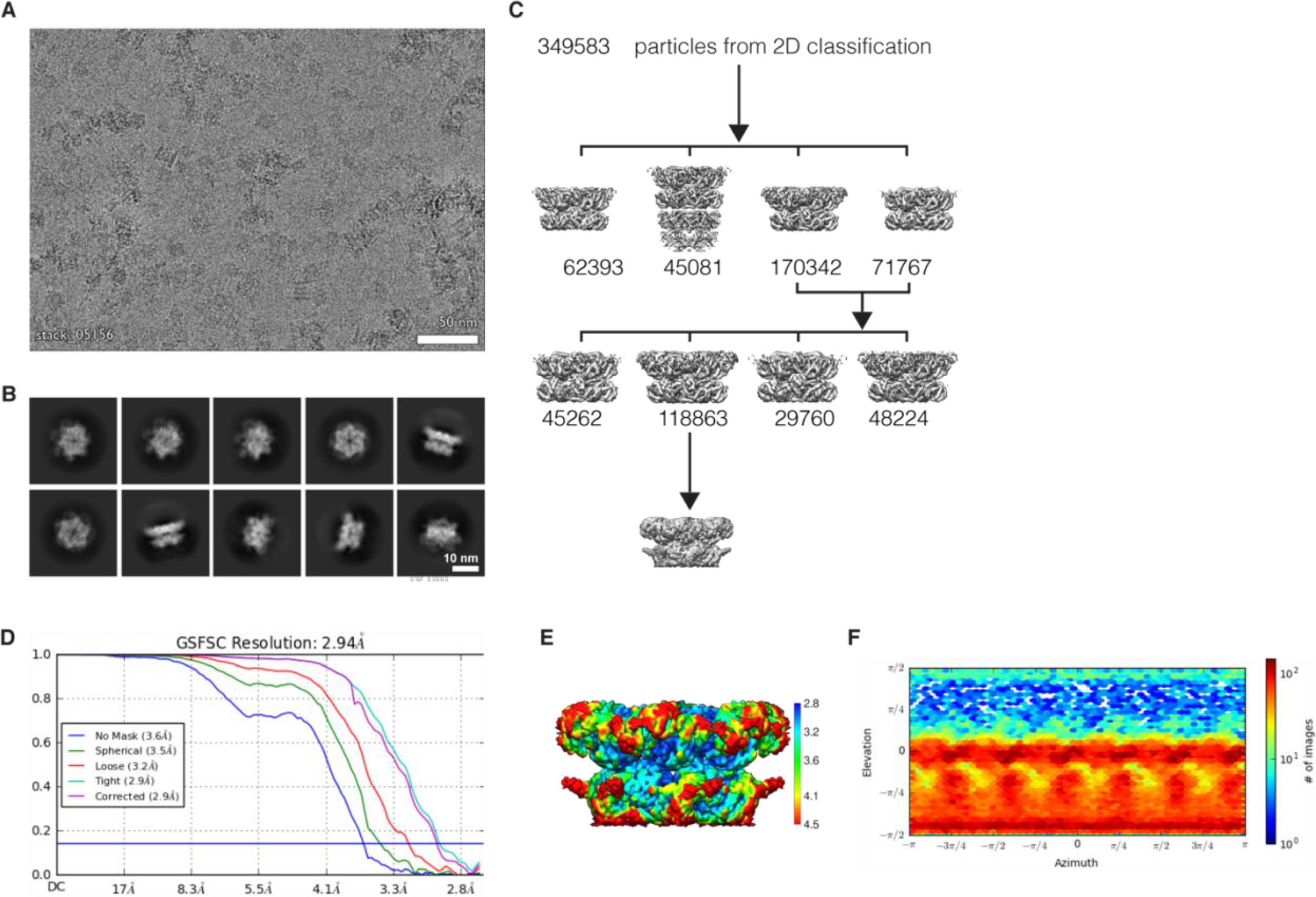
ADP-bound p97^D592N^. **(A)** A representative cryo-EM micrograph of ADP-bound p97^D592N^. **(B)** Representative 2D classes, showing multiple orientations. **(C)** Workflow of cryo-EM image processing. **(D)** FSC curves. **(E)** Cryo-EM side-view density of p97 mutants, colored according to local resolution and **(F)** Viewing direction distribution plot for mutant density.

**Fig. S14:**
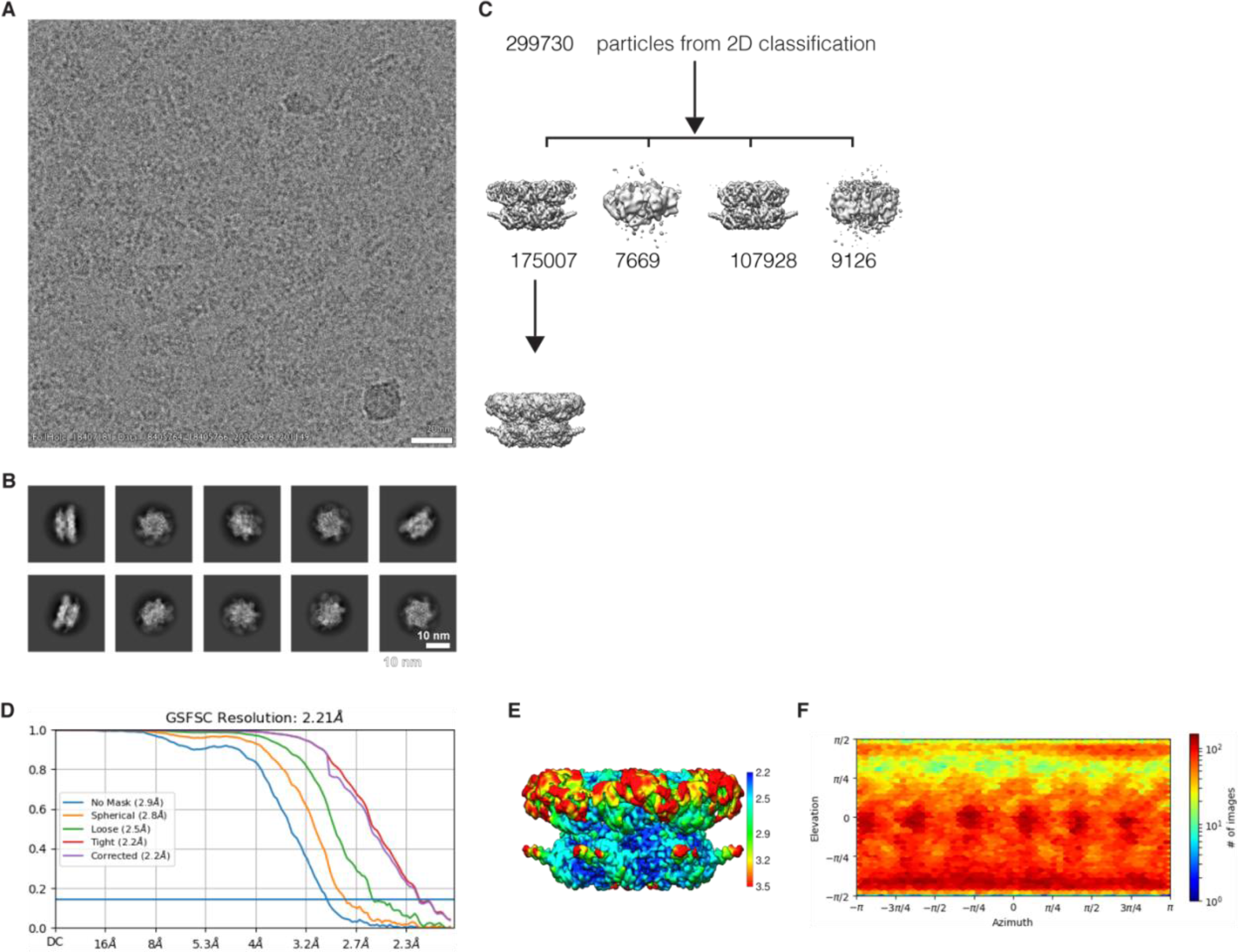
ATP*γ*S-bound p97^D592N^. **(A)** A representative cryo-EM micrograph of ATP*γ*S-bound p97^D592N^. **(B)** Representative 2D classes, showing multiple orientations. **(C)** Workflow of cryo-EM image processing. **(D)** FSC curves. **(E)** Cryo-EM side-view density of p97 mutants, colored according to local resolution and **(F)** Viewing direction distribution plot for mutant density.

**Fig. S15:**
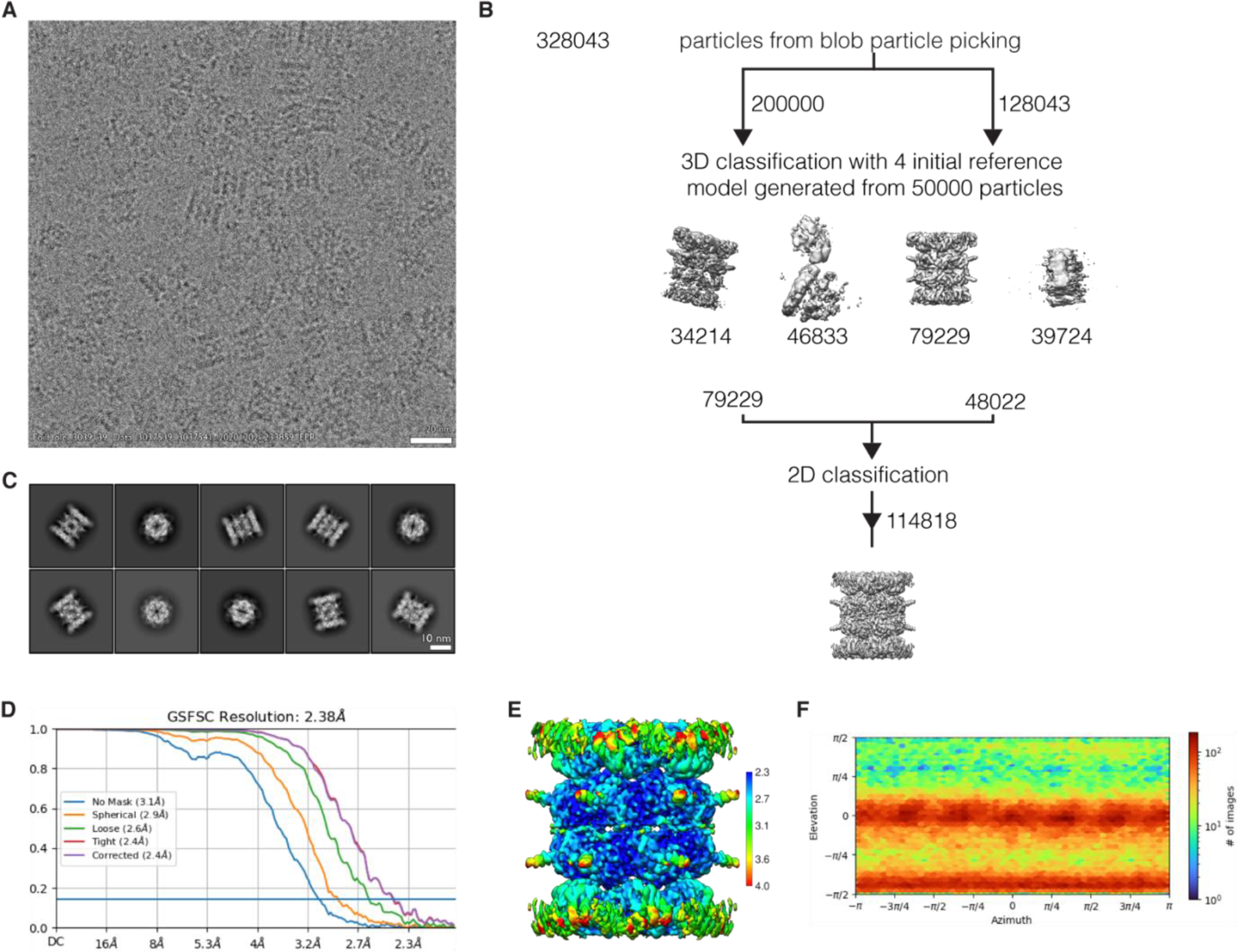
CB-5083-bound ADP-p97. **(A)** A representative cryo-EM micrograph of D1-ADP-D2-CB-5083-bound p97^WT^. **(B)** Representative 2D classes, showing multiple orientations. **(C)** Workflow of cryo-EM image processing. **(D)** FSC curves. **(E)** Cryo-EM side-view density of p97, colored according to local resolution and **(F)** Viewing direction distribution plot for mutant density.

**Fig. S16:**
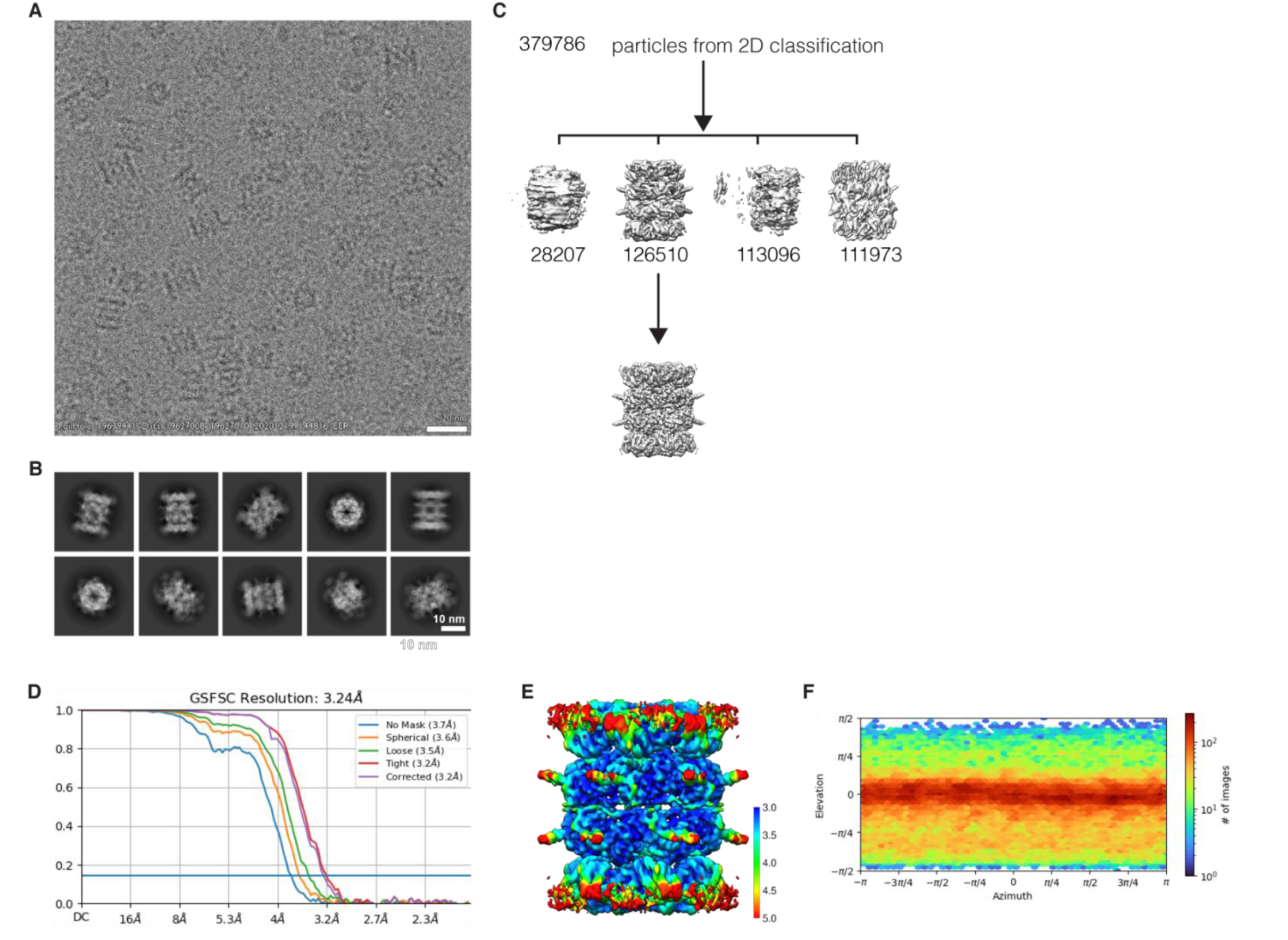
CB-5083-bound ATP*γ*S-p97. **(A)** A representative cryo-EM micrograph of D1-ATP*γ*S-D2-CB-5083-bound p97^WT^. **(B)** Representative 2D classes, showing multiple orientations. **(C)** Workflow of cryo-EM image processing. **(D)** FSC curves. **(E)** Cryo-EM side-view density of p97 mutants, colored according to local resolution and **(F)** Viewing direction distribution plot for mutant density.

## SUPPLEMENTAL TABLES

**Table S1.**
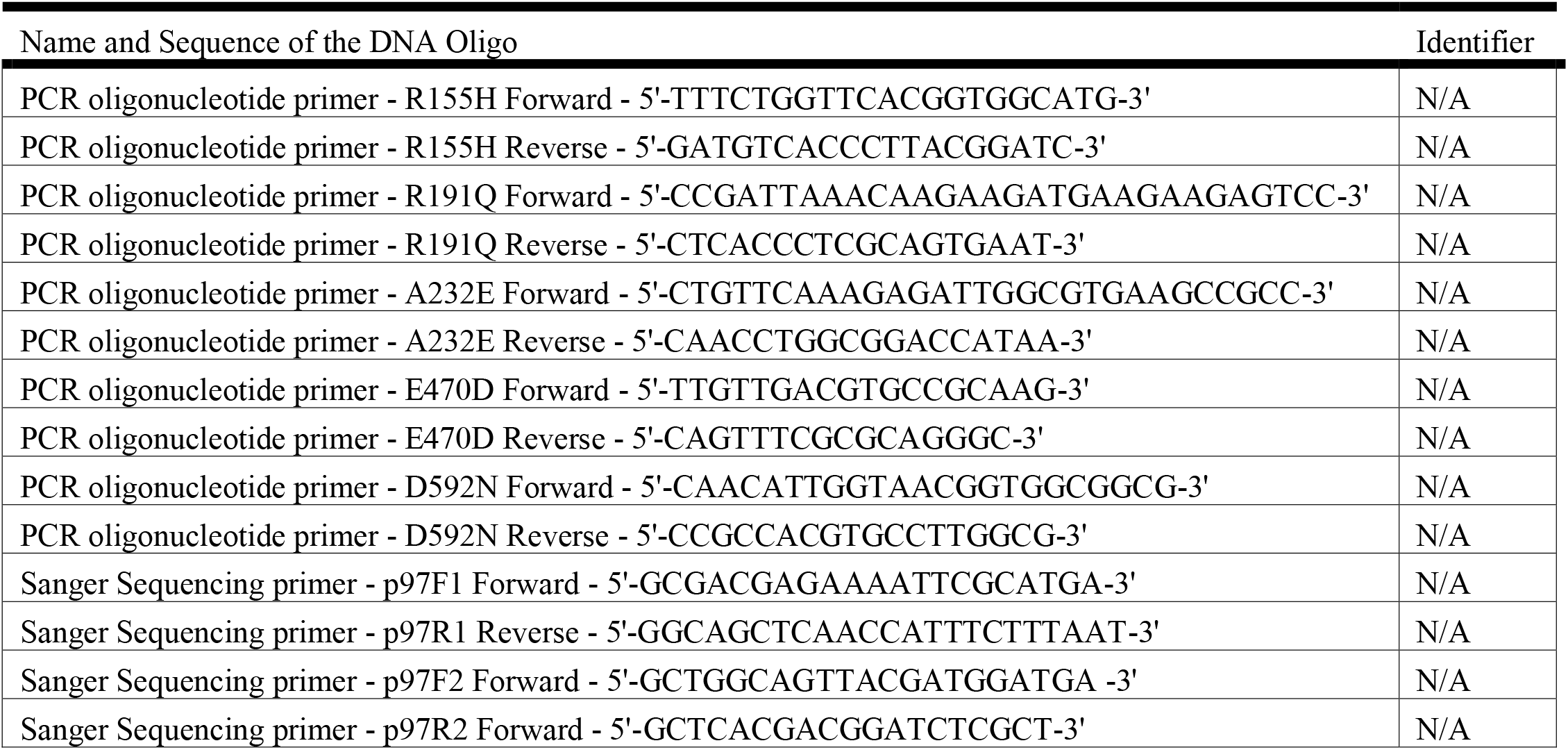
Oligonucleotides Synthesized by ThermoFisher for this Study, Related to Key Resources Table.

## REFERENCES

1. Erzberger, J. P., and Berger, J. M. (2006) Evolutionary relationships and structural mechanisms of AAA+ proteins. Annu. Rev. Biophys. Biomol. Struct. 35, 93–114

2. Stach, L., and Freemont, P. S. (2017) The AAA+ ATPase p97, a cellular multitool. Biochem. J. 474, 2953–2976

3. Cooney, I., Han, H., Stewart, M. G., Carson, R. H., Hansen, D. T., Iwasa, J. H., Price, J. C., Hill, C. P., and Shen, P. S. (2019) Structure of the Cdc48 segregase in the act of unfolding an authentic substrate. Science (80-.). 365, 502–505

4. Twomey, E. C., Ji, Z., Wales, T. E., Bodnar, N. O., Ficarro, S. B., Marto, J. A., Engen, J. R., and Rapoport, T. A. (2019) Substrate processing by the Cdc48 ATPase complex is initiated by ubiquitin unfolding. Science (80-.). 10.1126/science.aax1033

5. Banerjee, S., Bartesaghi, A., Merk, A., Rao, P., Bulfer, S. L., Yan, Y., Green, N., Mroczkowski, B., Neitz, R. J., Wipf, P., Falconieri, V., Deshaies, R. J., Milne, J. L. S., Huryn, D., Arkin, M., and Subramaniam, S. (2016) 2.3 Å resolution cryo-EM structure of human p97 and mechanism of allosteric inhibition. Science (80-.). 351, 871–875

6. Bulfer, S. L., Chou, T. F., and Arkin, M. R. (2016) P97 Disease Mutations Modulate Nucleotide-Induced Conformation to Alter Protein-Protein Interactions. ACS Chem. Biol. 11, 2112–2116

7. Deshaies, R. J. (2014) Proteotoxic crisis, the ubiquitin-proteasome system, and cancer therapy. BMC Med. 10.1186/s12915-014-0094-0

8. Roux, B., Vaganay, C., Vargas, J. D., Alexe, G., Benaksas, C., Pardieu, B., Fenouille, N., Ellegast, J. M., Malolepsza, E., Ling, F., Sodaro, G., Ross, L., Pikman, Y., Conway, A. S., Tang, Y., Wu, T., Anderson, D. J., Le Moigne, R., Zhou, H. J., Luciano, F., Hartigan, C. R., Galinsky, I., DeAngelo, D. J., Stone, R. M., Auberger, P., Schenone, M., Carr, S. A., Guirouilh-Barbat, J., Lopez, B., Khaled, M., Lage, K., Hermine, O., Hemann, M. T., Puissant, A., Stegmaier, K., and Benajiba, L. (2021) Targeting acute myeloid leukemia dependency on VCP-mediated DNA repair through a selective second-generation small-molecule inhibitor. Sci. Transl. Med. 10.1126/scitranslmed.abg1168

9. Arita, M., Wakita, T., and Shimizu, H. (2012) Valosin-Containing Protein (VCP/p97) Is Required for Poliovirus Replication and Is Involved in Cellular Protein Secretion Pathway in Poliovirus Infection. J. Virol. 86, 5541–5553

10. Huryn, D. M., Kornfilt, D. J. P., and Wipf, P. (2020) P97: An Emerging Target for Cancer, Neurodegenerative Diseases, and Viral Infections. J. Med. Chem. 63, 1892–1907

11. Watts, G. D. J., Wymer, J., Kovach, M. J., Mehta, S. G., Mumm, S., Darvish, D., Pestronk, A., Whyte, M. P., and Kimonis, V. E. (2004) Inclusion body myopathy associated with Paget disease of bone and frontotemporal dementia is caused by mutant valosin-containing protein. Nat. Genet. 36, 377–381

12. Niwa, H., Ewens, C. A., Tsang, C., Yeung, H. O., Zhang, X., and Freemont, P. S. (2012) The role of the N-domain in the atpase activity of the mammalian AAA ATPase p97/VCP. J. Biol. Chem. 287, 8561–8570

13. Blythe, E. E., Gates, S. N., Deshaies, R. J., and Martin, A. (2019) Multisystem Proteinopathy Mutations in VCP/p97 Increase NPLOC4·UFD1L Binding and Substrate Processing. Structure. 27, 1820–1829.e4

14. Johnson, J. O., Mandrioli, J., Benatar, M., Abramzon, Y., Van Deerlin, V. M., Trojanowski, J. Q., Gibbs, J. R., Brunetti, M., Gronka, S., Wuu, J., Ding, J., McCluskey, L., Martinez-Lage, M., Falcone, D., Hernandez, D. G., Arepalli, S., Chong, S., Schymick, J. C., Rothstein, J., Landi, F., Wang, Y. D., Calvo, A., Mora, G., Sabatelli, M., Monsurrò, M. R., Battistini, S., Salvi, F., Spataro, R., Sola, P., Borghero, G., Galassi, G., Scholz, S. W., Taylor, J. P., Restagno, G., Chiò, A., and Traynor, B. J. (2010) Exome Sequencing Reveals VCP Mutations as a Cause of Familial ALS. Neuron. 68, 857–864

15. Bastola, P., Wang, F., Schaich, M. A., Gan, T., Freudenthal, B. D., Chou, T. F., and Chien, J. (2017) Specific mutations in the D1-D2 linker region of VCP/p97 enhance ATPase activity and confer resistance to VCP inhibitors. Cell Death Discov. 10.1038/cddiscovery.2017.65

16. Tang, W. K., and Xia, D. (2013) Altered intersubunit communication is the molecular basis for functional defects of pathogenic p97 mutants. J. Biol. Chem. 288, 36624–36635

17. Schuetz, A. K., and Kay, L. E. (2016) A dynamic molecular basis for malfunction in disease mutants of p97/VCP. Elife. 10.7554/eLife.20143

18. Pan, M., Zheng, Q., Yu, Y., Ai, H., Xie, Y., Zeng, X., Wang, C., Liu, L., and Zhao, M. (2021) Seesaw conformations of Npl4 in the human p97 complex and the inhibitory mechanism of a disulfiram derivative. Nat. Commun. 10.1038/s41467-020-20359-x

19. Tang, W. K., Li, D., Li, C. C., Esser, L., Dai, R., Guo, L., and Xia, D. (2010) A novel ATP-dependent conformation in p97 N-D1 fragment revealed by crystal structures of disease-related mutants. EMBO J. 29, 2217–2229

20. Mills, J. E. J. (1996) Three-dimensional hydrogen-bond geometry and probability information from a crystal survey. J. Comput. Aided. Mol. Des. 10.1007/BF00134183

21. Shi, Z., Liu, S., Xiang, L., Wang, Y., Liu, M., Liu, S., Han, T., Zhou, Y., Wang, J., Cai, L., Gao, S., and Ji, Y. (2016) Frontotemporal dementia-related gene mutations in clinical dementia patients from a Chinese population. J. Hum. Genet. 61, 1003–1008

22. Hänzelmann, P., and Schindelin, H. (2017) The interplay of cofactor interactions and post-translational modifications in the regulation of the AAA+ ATPase p97. Front. Mol. Biosci. 10.3389/fmolb.2017.00021

23. Zhang, X., Gui, L., Zhang, X., Bulfer, S. L., Sanghez, V., Wong, D. E., Lee, Y. J., Lehmann, L., Lee, J. S., Shih, P. Y., Lin, H. J., Iacovino, M., Weihl, C. C., Arkin, M. R., Wang, Y., and Chou, T. F. (2015) Altered cofactor regulation with disease-associated p97/VCP mutations. Proc. Natl. Acad. Sci. U. S. A. 112, E1705–E1714

24. Zhou, H. J., Wang, J., Yao, B., Wong, S., Djakovic, S., Kumar, B., Rice, J., Valle, E., Soriano, F., Menon, M. K., Madriaga, A., Kiss Von Soly, S., Kumar, A., Parlati, F., Yakes, F. M., Shawver, L., Le Moigne, R., Anderson, D. J., Rolfe, M., and Wustrow, D. (2015) Discovery of a First-in-Class, Potent, Selective, and Orally Bioavailable Inhibitor of the p97 AAA ATPase (CB-5083). J. Med. Chem. 58, 9480–9497

25. Tang, W. K., Odzorig, T., Jin, W., and Xia, D. (2019) Structural basis of p97 inhibition by the site-selective anticancer compound CB-5083. Mol. Pharmacol. 95, 286–293

26. Rydzek, S., Shein, M., Bielytskyi, P., and Schütz, A. K. (2020) Observation of a Transient Reaction Intermediate Illuminates the Mechanochemical Cycle of the AAA-ATPase p97. J. Am. Chem. Soc. 143, 14472–1480

27. Punjani, A., Rubinstein, J. L., Fleet, D. J., and Brubaker, M. A. (2017) CryoSPARC: Algorithms for rapid unsupervised cryo-EM structure determination. Nat. Methods. 14, 290–296

28. Pettersen, E. F., Goddard, T. D., Huang, C. C., Couch, G. S., Greenblatt, D. M., Meng, E. C., and Ferrin, T. E. (2004) UCSF Chimera - A visualization system for exploratory research and analysis. J. Comput. Chem. 25, 1605–1612

29. Pettersen, E. F., Goddard, T. D., Huang, C. C., Meng, E. C., Couch, G. S., Croll, T. I., Morris, J. H., and Ferrin, T. E. (2021) UCSF ChimeraX: Structure visualization for researchers, educators, and developers. Protein Sci. 30, 70–82

30. Liebschner, D., Afonine, P. V., Baker, M. L., Bunkoczi, G., Chen, V. B., Croll, T. I., Hintze, B., Hung, L. W., Jain, S., McCoy, A. J., Moriarty, N. W., Oeffner, R. D., Poon, B. K., Prisant, M. G., Read, R. J., Richardson, J. S., Richardson, D. C., Sammito, M. D., Sobolev, O. V., Stockwell, D. H., Terwilliger, T. C., Urzhumtsev, A. G., Videau, L. L., Williams, C. J., and Adams, P. D. (2019) Macromolecular structure determination using X-rays, neutrons and electrons: Recent developments in Phenix. Acta Crystallogr. Sect. D Struct. Biol. 75, 861–877

31. Rose, R., Golosova, O., Sukhomlinov, D., Tiunov, A., and Prosperi, M. (2019) Flexible design of multiple metagenomics classification pipelines with ugene. Bioinformatics. 35, 1963–1965

32. Anderson, D. J., Le Moigne, R., Djakovic, S., Kumar, B., Rice, J., Wong, S., Wang, J., Yao, B., Valle, E., Kiss von Soly, S., Madriaga, A., Soriano, F., Menon, M. K., Wu, Z. Y., Kampmann, M., Chen, Y., Weissman, J. S., Aftab, B. T., Yakes, F. M., Shawver, L., Zhou, H. J., Wustrow, D., and Rolfe, M. (2015) Targeting the AAA ATPase p97 as an Approach to Treat Cancer through Disruption of Protein Homeostasis. Cancer Cell. 28, 653–665

33. Rosenthal, P. B., and Henderson, R. (2003) Optimal determination of particle orientation, absolute hand, and contrast loss in single-particle electron cryomicroscopy. J. Mol. Biol. 333, 721–745

